# Knockout of all nematode-specific NSPC genes expressed exclusively in the excretory gland cell results in transcriptomic signatures indicating an affected insulin signaling

**DOI:** 10.1101/2024.12.06.627198

**Authors:** Zuzanna Mackiewicz, Vladyslava Liudkovska, Andrzej Dziembowski

## Abstract

The nematode *Caenorhabditis elegans* is one of the best-studied model organisms in molecular biology; however, many aspects of its physiology and the functions of many genes remain poorly understood. In this study, we investigated the role of nematode-specific NSPC proteins, whose mRNAs were recently identified as primary targets of the poly(A) polymerase TENT-5. Surprisingly, we found that NSPCs are exclusively expressed in the excretory gland cell, a cell with still unclear functionality. Using an optogenetic approach, we precisely ablated the excretory gland cell and observed that nematodes exhibited no transcriptomic or physiological changes in its absence. Additionally, we generated and thoroughly studied a strain with a deletion of all 18 *nspc* genes, which revealed that, despite previous indications, NSPCs do not influence the worm’s defense response. Instead, the transcriptomic analysis showed that the absence of NSPCs strongly impacts DAF-2/DAF-16 insulin signaling, suggesting that NSPCs may function as neuropeptides influencing key *C. elegans* signaling pathways. Although further studies are required to elucidate the physiological effects of this regulation, our findings provide new insights into this unexplored part of nematode physiology.

## Introduction

The small nematode *Caenorhabditis elegans* was introduced to the scientific community in the 1970s and rapidly became one of the most important model organisms used in genetics and molecular biology [1]. Consequently, it is one of the best-studied multicellular organisms and was the first to have its entire genome sequenced [2], as well as its neuronal connectome [3,4] and cell lineage completely mapped [5–7]. Despite the extensive exploration of *C. elegans* over the years, some aspects of its anatomy and physiology remain poorly understood. Among these is the excretory gland cell, one of the 959 cells in the adult hermaphrodite worm, whose function remains a mystery, with only a few studies briefly mentioning its existence [8–11].

The excretory gland cell, along with the pore cell, duct cell, and excretory (canal) cell, constitutes the nematode’s secretory system, positioned on the ventral side of the pharynx, adjacent to the anterior section of the intestine [12]. This four-cell organ develops from the early embryonic stem cell (AB) and becomes functional at or soon after hatching. The excretory gland cell is binucleate and A-shaped and forms by the fusion of two identical cells at the 1.5-fold embryo stage. In addition to its tight connection with other excretory cells, it is also joined to the nerve ring, suggesting possible neural regulation of its activity. Structurally, the excretory gland cell features an extensive endoplasmic reticulum (ER), numerous mitochondria, and a dense ribosomal population [12].

In the 1980s, laser ablation was used to investigate the function and interactions of individual cells within the excretory system, including the excretory gland cell [10,11]. By precisely targeting cell nuclei with a laser beam at different developmental stages (L1, L2, young adult, or dauer), researchers observed that, while all other secretory cells are involved in osmoregulation, the excretory gland cell is not required for this process and also does not regulate *C. elegans* fertility or viability. Despite these findings made decades ago, no further studies have explored the function of the excretory gland cell.

In our previous research, we described the role of the non-canonical poly(A) polymerase TENT-5 in the innate immune response of *C. elegans* [13]. Using Nanopore Direct RNA sequencing, we demonstrated that TENT-5 polyadenylates and stabilizes a pool of mRNAs, many of which encode short, secreted proteins with well-defined or putative roles in the defense response. Intriguingly, our transcriptomic analysis revealed that the most prominent targets of TENT-5 were members of the NSPC family (Nematode Specific Peptide Family, group C), which encodes 18 small, highly similar proteins. Although some evidence suggests that NSPCs may be connected to immune or stress responses in worms [14–16], their exact function remains unknown. However, the observed relationship between TENT-5 and *nspc* mRNAs indicates that, like other TENT-5 targets, NSPC proteins may indeed be involved in immune regulation.

In this work, we show that all members of the NSPC family exhibit strong and highly specific expression exclusively in the excretory gland cell. This important finding prompted us to investigate the functions of both the excretory gland cell and NSPC proteins in *C. elegans* physiology. To uncover the role of the excretory gland cell, we employed an optogenetic approach to irreversibly damage the cell and disrupt its function. Since NSPCs are among the most abundant proteins in this cell, we also used a complementary approach by generating a CRISPR/Cas9 knockout strain lacking all *nspc* genes. Our extensive studies reveal that neither the ablation of the excretory gland cell nor the deletion of all NSPCs led to profound transcriptome changes or noticeable physiological phenotypes under tested laboratory conditions. However, here we propose that NSPCs might not possess antimicrobial properties, as predicted before, but might serve as neuropeptides that help regulate important signaling pathways such as the insulin DAF-2/DAF-16 pathway. Although our findings could not fully explain this mechanism, they provide valuable insights into this unexplored aspect of nematode biology, and we believe they will accelerate future research in the field.

## Results and discussion

### NSPCs are exclusively expressed in the excretory gland cell

The NSPC family comprises 18 genes clustered on chromosome X (Figure 1A), encoding small, secreted proteins with highly similar sequences and structures (Figure 1B and C) that are not related to any other known proteins outside the *Caenorhabditis* genus. Surprisingly, only one study has explored the possible functions of NSPC members, proposing that they may have antimicrobial activity based on their sequence and structure [14]. This prediction is consistent with high-throughput transcriptomic analyses showing dysregulation of *nspc* genes mostly upon various pathogen infections (Supplementary Table S1). For example, bacterial infection with *Enterococcus faecalis* (OG1RF), *Photorhabdus luminescens* (HB), or *Xenorhabdus nematophila* leads to the downregulation of multiple *nspc* genes [15,16]. In contrast, bacterial infection with *Pseudomonas aeruginosa* or *Serratia marcescens*, as well as fungal infection with *Drechmeria coniospora* or *Harposporium sp.,* results in increased expression levels of *nspc* genes [15–17]. Moreover, our previous studies on poly(A) polymerase TENT-5 function in *C. elegans* physiology revealed its pronounced activity toward NSPC-coding mRNAs, along with multiple known immune effectors [13]. However, despite many indications, NSPC activity has never been demonstrated either *in vitro* or *in vivo,* and their function remains unknown.

**Figure 1.**
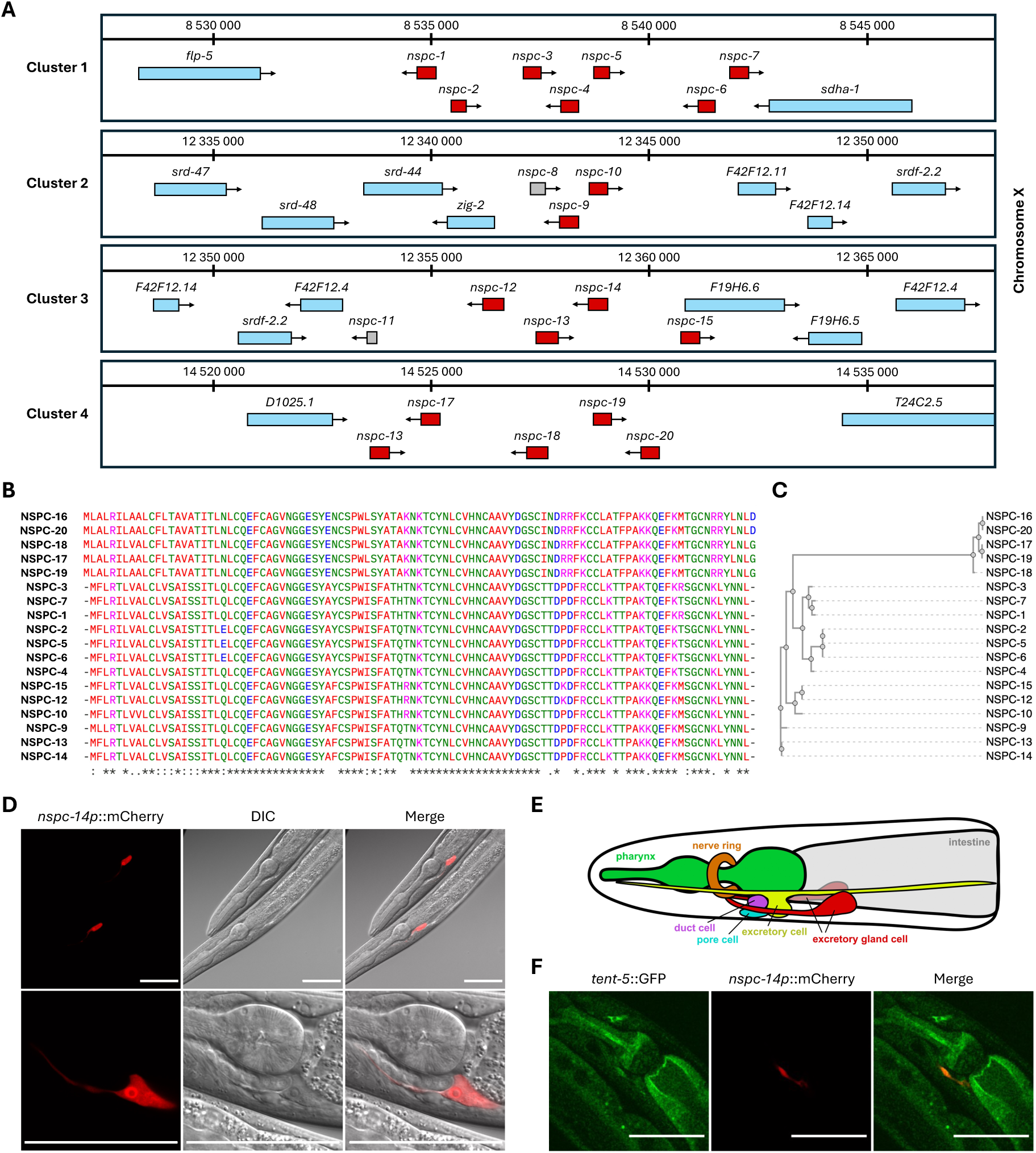
NSPCs are exclusively expressed in the excretory gland cell. **(A)** Genomic locations of *nspc* genes on chromosome X. *Nspc* genes are marked in red, neighboring genes in blue, and pseudogenes *nspc-8* and *nspc-11* in gray. Boxes represent gene structures, including exons, introns, and UTRs, while arrows indicate the DNA strand. The genomic map was prepared based on the WormBase genome browser [19]. **(B)** Multiple sequence alignment of NSPC proteins. Asterisks (*) represent residues conserved across all NSPCs; colons (:) indicate moderately conserved residues; and dots (.) indicate non-conserved residues with similar chemical properties. Alignment was performed using the MUSCLE tool from EMBL-EBI [20]. **(C)** Phylogenetic tree of NSPC proteins. Analysis was performed using the MUSCLE tool from EMBL-EBI [20]. **(D)** Fluorescence and differential interference contrast (DIC) microscopy images of NSPC-mCherry expression in *C. elegans* excretory gland cell. Scale bars => 50 µm. **(E)** Schematic representation of the excretory system in *C. elegans*. Prepared based on the scheme from Sundaram M.V. and Buechner M. (2016) [8]. The excretory system is located near the pharynx (green) and nerve ring (orange) and consists of the excretory cell (yellow), duct cell (purple), pore cell (blue), and excretory gland cell (red). **(F)** Fluorescence microscopy images showing colocalization of TENT-5-GFP and NSPC-mCherry in the excretory gland cell. Scale bars => 50 µm.

Therefore, we aimed to study the exact role of NSPCs and investigate the importance of their relationship with TENT-5. We began by studying the localization of NSPCs within the *C. elegans* body. We generated extrachromosomal reporters for three *nspc* genes fused with mCherry and examined their expression using confocal microscopy. Surprisingly, we observed that NSPCs are expressed exclusively in a single cell – the excretory gland cell (Figure 1D, Supplementary Figure S1), which agrees with high-throughput cell-specific expression profiling [18]. Moreover, the expression levels of NSPCs are extremely high, as they are detectable in RNA sequencing experiments, even from whole-animal samples. Although not all NSPCs exhibit the same expression levels throughout *C. elegans’s* lifespan, they appear to be highly expressed throughout all developmental stages [18,19]. The excretory gland cell is a part of the nematode’s excretory system, where four distinct cells collaborate to perform a specific task throughout the nematode’s lifespan (Figure 1E). Notably, while the functions of the other excretory cells have been characterized, the role of the excretory gland cell, similarly to NSPC proteins, remains a mystery.

We also checked whether NSPCs and TENT-5 colocalize in the excretory gland cell using worms expressing both *tent-5*::GFP and *nspc-14p*::mCherry as extrachromosomal arrays. Indeed, we found that TENT-5 is also expressed in the excretory gland cell (Figure 1F), allowing for the direct regulation of *nspc* mRNAs by TENT-5. Following these discoveries, here we aimed to explain the function of the excretory gland cell, understand why it requires such a high and specific expression of NSPCs, and explore the role of TENT-5 in regulating this previously undescribed aspect of *C. elegans* physiology.

### Optogenetic ablation can be successfully implemented to destroy the excretory gland cell

To study the role of the excretory gland cell, we took advantage of the restricted NSPC expression, enabling the optogenetic cell ablation approach, which has been previously used for the precise ablation of neurons, muscles, and epidermal cells in *C. elegans* [21,22]. This method relies on the expression of the genetically encoded photosensitizer miniSOG (mini Singlet Oxygen Generator) specifically in the cell or tissue of interest [23]. Upon blue light illumination (maximum excitation at 448 nm), miniSOG is targeted to the outer mitochondrial membrane, where it generates toxic levels of singlet oxygen (^1^O_2_) and other reactive oxygen species (ROS), leading to cell damage and death (Figure 2A).

**Figure 2.**
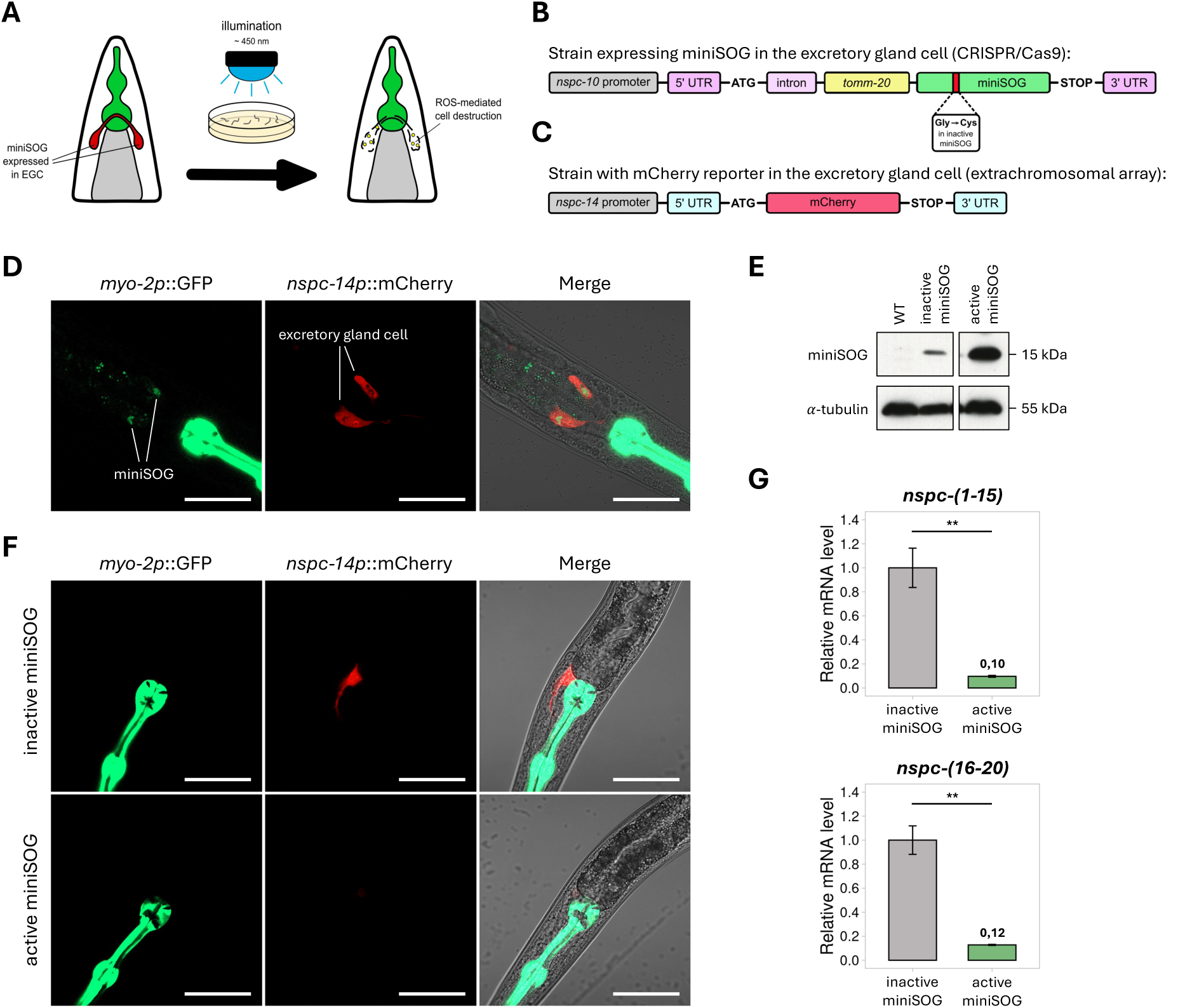
Optogenetic ablation can be implemented to destroy the excretory gland cell. **(A)** Schematic representation of the optogenetic ablation of the excretory gland cell (EGC). Worms were illuminated on plates using the LED board emitting 460 nm blue light in five 10-minute intervals. Upon illumination, miniSOG generates reactive oxygen species (ROS), leading to cell-specific destruction. **(B)** Scheme illustrating the design of *C. elegans* transgenic strains used for the optogenetic ablation of the excretory gland cell. miniSOG, together with mitochondria-directing TOMM-20, was expressed under the *nspc-10* promoter, with the *nspc-10* intron and 5/3ʹ UTRs. A single nucleotide mutation in the miniSOG sequence results in Gly->Cys amino acid substitution, blocking miniSOG activity. **(C)** Scheme showing the design of *C. elegans* transgenic strain used for visualization of the excretory gland cell. Red fluorescence protein mCherry was expressed under the *nspc-14* promoter in the extrachromosomal array. **(D)** Fluorescence microscopy images of the excretory gland cell expressing miniSOG. Green fluorescence of miniSOG is visible in the cytoplasm of the excretory gland cell, likely overlapping with mitochondria localization. Scale bars => 50 µm. **(E)** Western blot showing successful generation of two miniSOG-expressing strains. α-tubulin was used as a loading control. The lower amount of miniSOG in the inactive miniSOG strain may result from suboptimal antibody binding to the mutated protein. **(F)** Fluorescence microscopy images of the excretory gland cell after optogenetic ablation. Top images show a worm with inactive miniSOG (control), and the bottom – a worm with active miniSOG. The absence of red NSPC-mCherry fluorescence in the worm with active miniSOG indicates successful ablation. Scale bars => 50 µm. **(G)** RT-qPCR showing a decrease in all *nspc* gene expression levels after ablation of the excretory gland cell. Relative *nspc* mRNA levels were normalized to *act-1*. Bar plots represent mean values with SD. ** indicates *p*-value < 0.01 (two-tailed *t-*test).

Using CRISPR/Cas9, we generated two transgenic strains expressing miniSOG under the *nspc-10* promoter: one with active miniSOG fused to the mitochondrial membrane protein TOMM-20 and another with miniSOG carrying a point mutation that renders the photosensitizer inactive, serving as a negative control (Figure 2B). To visualize the excretory gland cell and monitor its damage post-ablation, both miniSOG strains were crossed with a strain expressing the red-fluorescent protein mCherry under the *nspc-14* promoter on the extrachromosomal array (Figure 2C). Since miniSOG itself is a green fluorescent flavoprotein [23], we observed its localization to the excretory gland cell (Figure 2D) and also confirmed protein production by western blot analysis (Figure 2E), thus validating the correct strain preparation for optogenetic ablation.

To optimize the ablation protocol for the excretory gland cell, we tested various blue light sources, illumination durations, and developmental stages of the worms (Supplementary Figure S2A). Unlike other optogenetic studies, where photosensitizers were typically activated using microscope fluorescence lamps, we developed a high-throughput method using an LED advertising board emitting 460 nm blue light (Supplementary Figure S2B). This setup enabled the simultaneous processing of a large worm population for subsequent high-throughput analyses, such as RNA sequencing. Ultimately, we achieved optimal excretory gland cell ablation by illuminating L3 stage larvae with continuous blue light in five 10-minute intervals, allowing time to prevent worm overheating. This approach successfully and selectively destroyed the excretory gland cell in worms expressing active miniSOG, while worms expressing the mutated control showed no signs of cell damage. The ablation was confirmed by the loss of mCherry fluorescence approximately 40 hours post-illumination (Figure 2F, Supplementary Figure S2C) and by RT-qPCR, which demonstrated a nearly 90% reduction in NSPCs expression (Figure 2G). These observations indicate that the cell was either efficiently destroyed or significantly damaged, impairing its function.

In conclusion, we successfully employed a miniSOG-mediated optogenetic approach to ablate the excretory gland cell in *C. elegans*. This method provides a model for studying the function of the excretory gland cell and offers extensive possibilities for further investigation, surpassing the limitations of traditional laser ablation techniques.

### Ablation of the excretory gland cell leads to no transcriptome changes

We performed the optogenetic ablation of the excretory gland cell, as described above, and monitored worms for potential abnormal phenotypes. Interestingly, worms remained viable after ablation and displayed no visible phenotypes, which is consistent with results from previous laser ablation experiments [10,11]. A few possibilities might explain this surprising outcome. One possibility is that the excretory gland cell plays a more significant role during earlier developmental stages, which are challenging to study using the optogenetic setup due to the harmful effects of blue light exposure on younger worms (Supplementary Figure S2A). Another possibility is that defects in the excretory gland cell only become evident under some specific stress conditions. Furthermore, the loss of the excretory gland cell might lead to subtle effects that are not apparent in anatomical or physiological phenotypes but may instead be revealed through altered patterns of gene expression. To explore this latter possibility, we performed Illumina RNA sequencing (RNA-seq) on adult worms with both active and inactive (control) miniSOG two days after illumination. The RNA-seq results showed significant changes in gene expression, with 1210 genes downregulated and 1293 upregulated following cell ablation (Figure 3A, Supplementary Table S2). Notably, the downregulation of nearly all NSPC genes confirmed the successful removal and dysfunction of the excretory gland cell.

**Figure 3.**
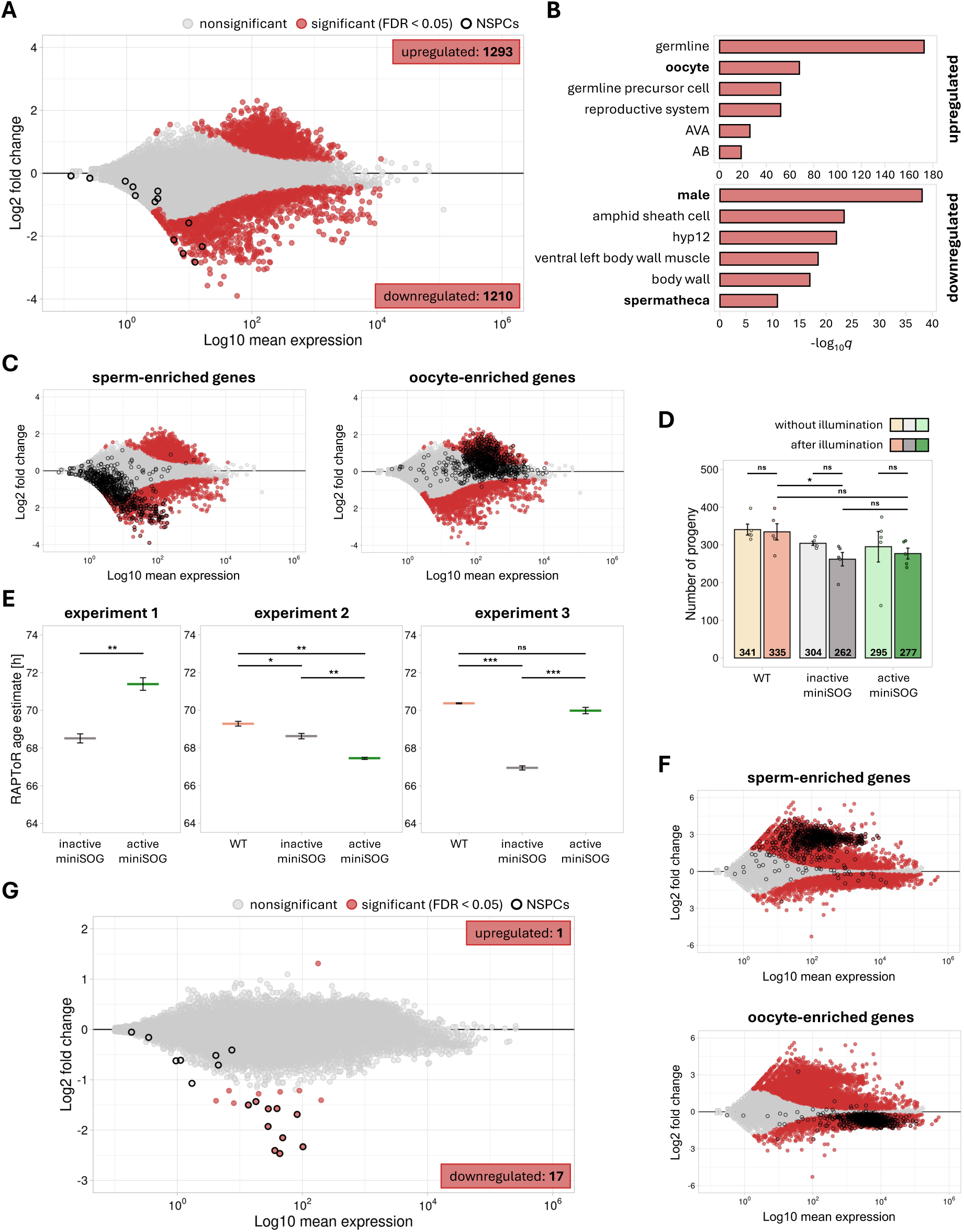
Ablation of the excretory gland cell leads to minimal transcriptome changes. **(A)** MA plot illustrating differential gene expression between inactive and active miniSOG worms after the successful ablation of the excretory gland cell. Significantly changed genes (FDR < 0.05) are marked with red dots. The black borderline highlights *nspc* genes. **(B)** Top GO terms for the upregulated (top) and downregulated (bottom) genes following the ablation of the excretory gland cell ordered by adjusted *p*-value. Terms related to the dysregulation of oocyte- and sperm-enriched genes are shown in bold. **(C)** Duplicates of the MA plot in **(A)** showing the downregulation of sperm-enriched genes and the upregulation of oocyte-enriched genes. Marked genes are listed in Supplementary Table S2 and are derived from Reinke V. *et al.* (2004) [28]. **(D)** Differences in brood sizes between wild-type (orange), inactive (gray), and active miniSOG (green) worms without and after the illumination (lighter and darker shades, respectively). Bar plots represent mean values with SD (n=5). ns => not significant; * => *p*-value < 0.05 (two-tailed *t-*test). **(E)** Age estimates for wild-type (orange), inactive (gray), and active miniSOG (green) worms after illumination, based on three independent RNA-seq replicates. Plots represent mean values with SD. ns => not significant; * => *p*-value < 0.05; ** => *p*-value < 0.01; *** => *p*-value < 0.001 (two-tailed *t-*test). **(F)** MA plots showing differential gene expression between wild-type and inactive miniSOG worms in the third RNA-seq experiment (two control conditions). Significantly changed genes (FDR < 0.05) are marked with red dots. Black borderline is used to mark sperm- and oocyte-enriched genes, as in **(C)**. **(G)** MA plots showing differential gene expression between inactive and active miniSOG worms after excretory gland cell ablation averaged across three independent RNA-seq experiments. Significantly changed genes (FDR < 0.05) are marked with red dots. Black borderline is used to mark *nspc* genes.

Interestingly, further analysis revealed that mainly sperm-enriched genes were downregulated, while oocyte-enriched genes were upregulated (Figure 3B and C). Although this pattern might suggest a role for the excretory gland cell in reproduction, we did not detect any impact of this gene dysregulation on worm brood sizes following ablation (Figure 3D). However, the observed transcriptome changes might also reflect a characteristic gene expression profile associated with age differences between worms, since sperm gene expression peaks in the L4 stage, while oocyte genes are expressed more in young adults [24,25]. To test this possibility, we used the R package RAPToR [26] to estimate the age of the worms in each RNA-seq sample (Figure 3E). Indeed, although all samples were processed simultaneously and appeared to be of similar age based on microscopic observations, we found an age difference of a few hours between worms with active and inactive miniSOG, which likely contributed to the overall gene expression pattern.

To minimize confusion caused by age-related differences, we repeated the RNA sequencing two more times and included wild-type worms post-illumination as an additional control. Unexpectedly, the transcriptome changes remained variable but consistently involved sperm- and oocyte-enriched genes (Supplementary Figure S3), indicating developmental stage variations between samples (Figure 3E). These differences were also strongly pronounced between two control samples in one of the experiments (Figure 3E and F, Supplementary Table S2), supporting the idea that age discrepancies influenced the results. The exact reason for such age variance between conditions in this particular experiment remains unclear, but we suspect that the illumination process may have impacted the worms’ developmental timing [27], even though no changes in behavior or viability were observed afterward (Supplementary Figure S3).

Finally, to reduce the impact of the observed age differences, we combined the results from all three independent experiments. Surprisingly, the overall transcriptome changes following excretory gland cell ablation were minimal, with only 17 genes significantly downregulated and one upregulated (Figure 3G, Supplementary Table S2). Among the downregulated genes, ten belonged to the NSPC family, confirming the successful ablation of the excretory gland cell but also suggesting that the cell’s function may primarily involve NSPC production.

The minimal impact on both nematode physiology and transcriptome following the ablation of the entire cell seems quite unusual. As mentioned above, the cell’s role may be manifested under the environmental pressures found in the wild, but it is difficult to detect in standard laboratory conditions. More extensive studies will be required to further explore this hypothesis.

### NSPC deletion results in no detectable physiological phenotypes

The averaged RNA sequencing results following the excretory gland cell ablation suggest that its primary role is likely connected with NSPC production. However, the functions of NSPC proteins remain largely unknown, complicating the interpretation of the optogenetic ablation results and making it challenging to assess the cell’s overall significance. Therefore, we aimed to clarify the function of this poorly understood gene family, explain the strong TENT-5-mediated regulation of NSPCs, and provide new insights into the possible role of the excretory gland cell.

Given the similarity within NSPCs (Figure 1B and C), we initially assumed that RNA interference (RNAi), a method frequently and successfully employed in *C. elegans* research [29], would effectively silence all family members simultaneously. After extensive optimization, we achieved up to 90% silencing of NSPCs, followed by extensive transcriptomic and physiological experiments (Supplementary Figure S4A-D, Supplementary Table S3). However, due to the variability in silencing 18 genes simultaneously, RNAi results were often inconsistent, which complicated result interpretation. To ensure data reliability, we generated a CRISPR/Cas9 knockout strain lacking all NSPC genes. This process involved sequential deletions of four gene clusters: *nspc-1* – *nspc-7*, *nspc-8* – *nspc-10*, *nspc-11* – *nspc-15,* and *nspc-16* – *nspc-20* (Figure 4A).

**Figure 4.**
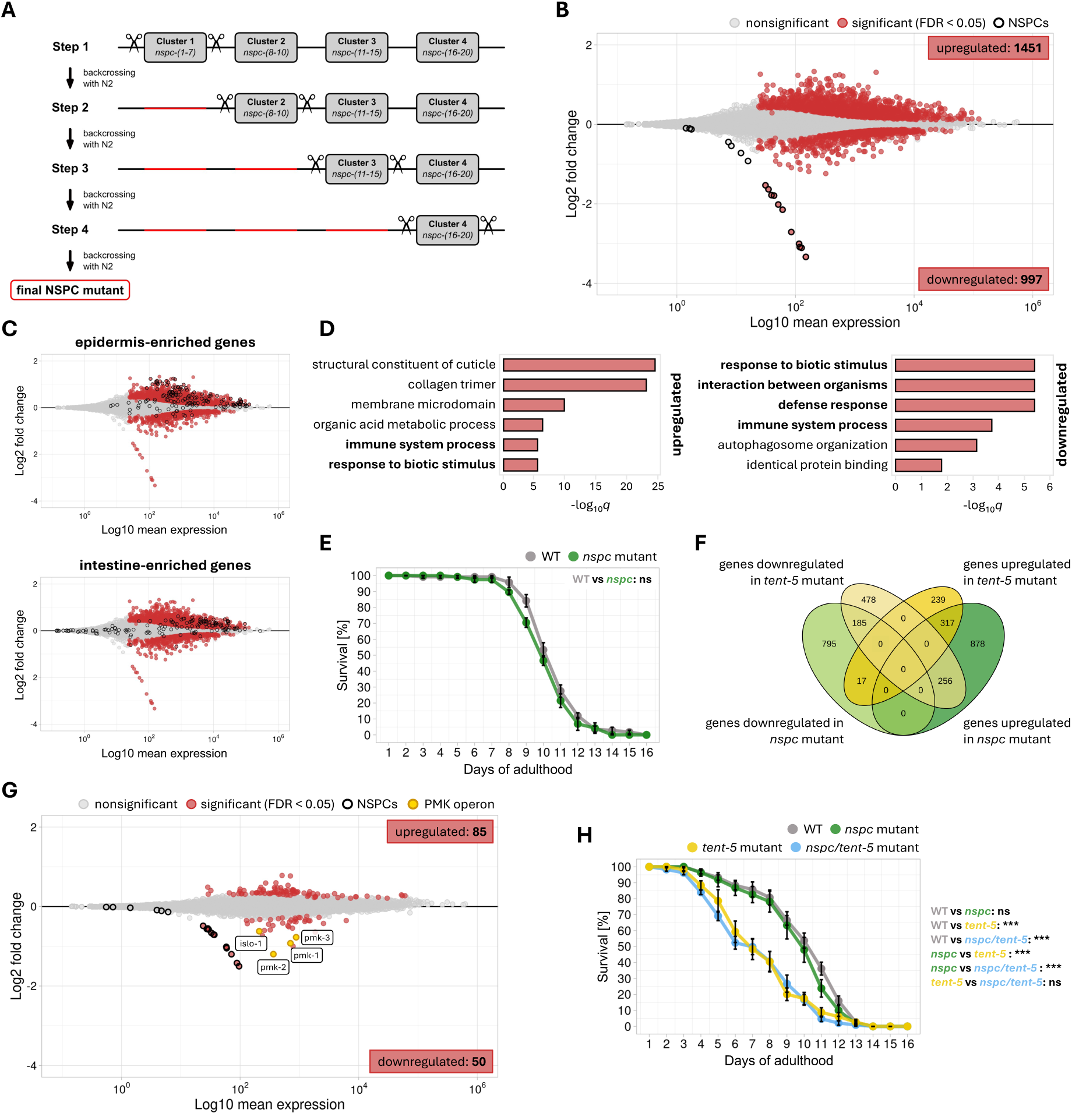
NSPC deletion does not result in any physiological phenotypes. **(A)** Scheme illustrating the design and generation of the null *nspc* mutant. Clusters depicted in Figure 1A were subsequently deleted using CRISPR/Cas9 technology. After each deletion, worms were outcrossed with wild-type males. **(B)** MA plots showing differential gene expression between wild-type and *nspc* mutant worms. Significantly changed genes (FDR < 0.05) are marked with red dots. Black borderline highlights *nspc* genes. **(C)** Duplicates of MA plot in **(B)** showing the upregulation of epidermis- and intestine-enriched genes in *nspc* mutant worms. Marked genes are listed in Supplementary Table S3 and are derived from CeNGEN database [18,38]. **(D)** Top GO terms for the 200 most upregulated (left) and downregulated (right) genes in the *nspc* mutant, ordered by adjusted *p*-value. Terms related to the defense response are shown in bold. **(E)** Survival of wild-type (gray) and *nspc* mutant (green) worms grown on non-pathogenic *E. coli* HB101. Points represent mean values from 6 separate plates (n≅20 each) with SD. ns => not significant (log-rank test). **(F)** Venn diagram showing overlaps between upregulated (darker shade) and downregulated (lighter shade) genes in *tent-5* (yellow) and *nspc* mutant (green) worms. Included are all significantly changed genes (FDR < 0.05). **(G)** MA plots showing differential gene expression between *tent-5* and *nscp/tent-5* mutant worms. Significantly changed genes (FDR < 0.05) are marked with red dots. Black borderline marks *nspc* genes. Yellow dots with an orange borderline show significantly downregulated genes from a single operon: *islo-1*, *pmk-1*, *pmk-2*, and *pmk-3*. **(H)** Survival of wild-type (gray), *nspc* mutant (green), *tent-5* mutant (yellow), and *nspc/tent-5* mutant (blue) worms grown on pathogenic *P. aeruginosa* PAO1. Points represent mean values from 6 separate plates (n≅20 each) with SD. ns => not significant; *** => *p*-value < 0.001 (log-rank test).

First, we performed RNA-seq and identified significant changes in gene expression following NSPC deletion, with 1451 genes upregulated and 997 downregulated compared to wild-type worms (Figure 4B, Supplementary Table S3). Interestingly, many upregulated genes showed tissue enrichment in the worm’s epidermis and intestine (Figure 4C), which are the first tissues to contact pathogens during infections and initiate immune response. Consistently with functional enrichment analysis, among the upregulated genes, we identified multiple regulators of innate immunity, such as infection response genes (4 *irg* family members) [30,31], C-type lectins (24 *clec* family members) [32], lysozymes (4 *lys* family members) [33], or aspartyl proteases (8 *asp* family members) [34] (Figure 4D). Similar GO terms were also observed for downregulated genes, suggesting a potential role for NSPCs in *C. elegans* defense responses. However, the observed expression changes, although significant, were generally small, with only a few genes exceeding a two-fold change. This modest gene expression change is consistent with the absence of visible phenotypes or significant lifespan alterations in the mutant worms (Figure 4E).

The potential involvement of NSPCs in defense processes, as indicated by our RNA-seq results, aligns with evolutionary predictions made for those genes [14] and the previously observed relationship between NSPCs and TENT-5 [13]. We previously demonstrated that TENT-5 stabilizes NSPC transcripts by elongating their poly(A) tails, although the role of this co-regulation in mediating the *C. elegans* response to pathogens remains unexplored. To better understand this mechanism, we first compared the pool of transcripts that were up- or downregulated following NSPC or TENT-5 deletion and found no clear overlaps between the groups, with some genes showing opposite regulation in the two mutants (Figure 4F, Supplementary Table S3). Next, using Nanopore Direct RNA Sequencing (DRS), we measured poly(A) tail lengths in *nspc* mutant worms and observed almost no changes in poly(A) metabolism compared to wild-type worms (Supplementary Figure S4E, Supplementary Table S4). This observation allowed us to exclude the possibility that TENT-5 is regulated or regulates other transcripts via NSPCs. Moreover, we generated a mutant strain lacking both TENT-5 and NSPCs and conducted additional RNA sequencing to analyze dependencies between these proteins (Figure 4G, Supplementary Figure S4F, Supplementary Table S3 and S4). Interestingly, NSPC deletion in the absence of TENT-5 led to smaller gene expression changes than when TENT-5 was present (Figure 4G, Supplementary Table S3), confirming that TENT-5 indeed regulates NSPCs. Among the few downregulated genes, we identified a unique operon containing the genes *islo-1, pmk-1, pmk-2,* and *pmk-3*, three of which are members of the PMK family, known for its crucial role in regulating *C. elegans* defense responses [35–37]. We showed that this result is due to the strong enrichment of PMK members in *tent-5* mutant worms, with levels returning to normal upon additional NSPC deletion (Supplementary Figure S4G), hinting at a possible complex interplay between TENT-5, NSPCs, and PMK pathway in regulating defense mechanisms.

Together, our results show that NSPCs are not essential for viability; however, they may play a role in facilitating defense responses. To further investigate this hypothesis, we conducted lifespan experiments with *P. aeruginosa* PAO1 infection, comparing four different *C. elegans* strains: wild-type N2, *tent-5* mutant, *nspc* mutant, and double *nspc*/*tent-5* mutant (Figure 4H). Although we confirmed previous findings of increased susceptibility of *tent-5* mutant worms to bacterial infection [13], we demonstrated that, despite previous indications, NSPCs are not required for worms’ defense response to *P. aeruginosa*. Additionally, our results suggest that the excretory gland cell, which is the hub for the NSPCs production, is also unlikely to play a direct role in innate immunity. However, based on our results, we cannot rule out the possibility that the excretory gland cell and NSPCs may be involved in responses to other pathogens.

### NSPCs and TENT-5 might be involved in regulating cholesterol metabolism

Extensive polyadenylation of *nspc* mRNAs by TENT-5 and the strong, cell-specific expression of NSPCs in the excretory gland cell indicate a functional link between TENT-5 and NSPCs, suggesting their potential importance for excretory gland cell activity. Unfortunately, our research indicates that while TENT-5 plays a crucial role in the *C. elegans* defense response, NSPCs do not, challenging earlier assumptions about their antimicrobial activity.

However, in our previous work, we speculated that TENT-5 might also be involved in regulating cholesterol metabolism [39]. In *C. elegans* males, cholesterol levels are significantly higher in male-specific tissues [40], where TENT-5 activity is strongest. Interestingly, cholesterol is also abundant in the excretory gland cell [40], leading us to hypothesize that TENT-5 activity may be exaggerated in cholesterol-rich tissues, influencing cholesterol metabolism in worms. Additionally, cholesterol is known to influence the p38/PMK-1 MAPK pathway [41,42], which is critical for innate immune responses in worms, further linking this hypothesis to our previous observation that PMK genes are significantly upregulated in *tent-5* mutants. Given that TENT-5 positively regulates the expression of NSPCs, we hypothesized that NSPCs could also be involved in cholesterol regulation. To explore this possibility, we conducted lifespan experiments on non-pathogenic *E. coli* and pathogenic *P. aeruginosa* for four *C. elegans* strains (wild type, *tent-5*, *nspc*, and *nspc*/*tent-5* mutants) under standard conditions and with varying cholesterol concentrations (Figure 5A-D). As expected, lower cholesterol levels generally reduced worm lifespan under both conditions (Figure 5A and B). Interestingly, when fed on non-pathogenic bacteria, *nspc* mutants performed slightly better than other strains at low cholesterol levels (Figure 5C), suggesting that NSPCs might negatively regulate cholesterol uptake or metabolism. Interestingly, *tent-5* mutants showed a shortened lifespan on both low and normal cholesterol plates (Figure 5A), indicating that TENT-5 might be required to facilitate some cholesterol-related processes. Finally, when both *tent-5* and *nspc* genes were deleted, the opposing effects of these proteins mitigated the impact of cholesterol on nematode lifespan (Figure 5A). However, these trends were not observed under pathogenic conditions, possibly because the effects of infection outweighed cholesterol-related changes (Figure 5B and D). Nonetheless, the difference between wild-type and *nspc* mutants compared to *tent-5* and *nspc*/*tent-5* mutants was slightly smaller after exposure to high cholesterol, suggesting that cholesterol might contribute to pathogen resistance in *C. elegans* (Figure 5D).

**Figure 5.**
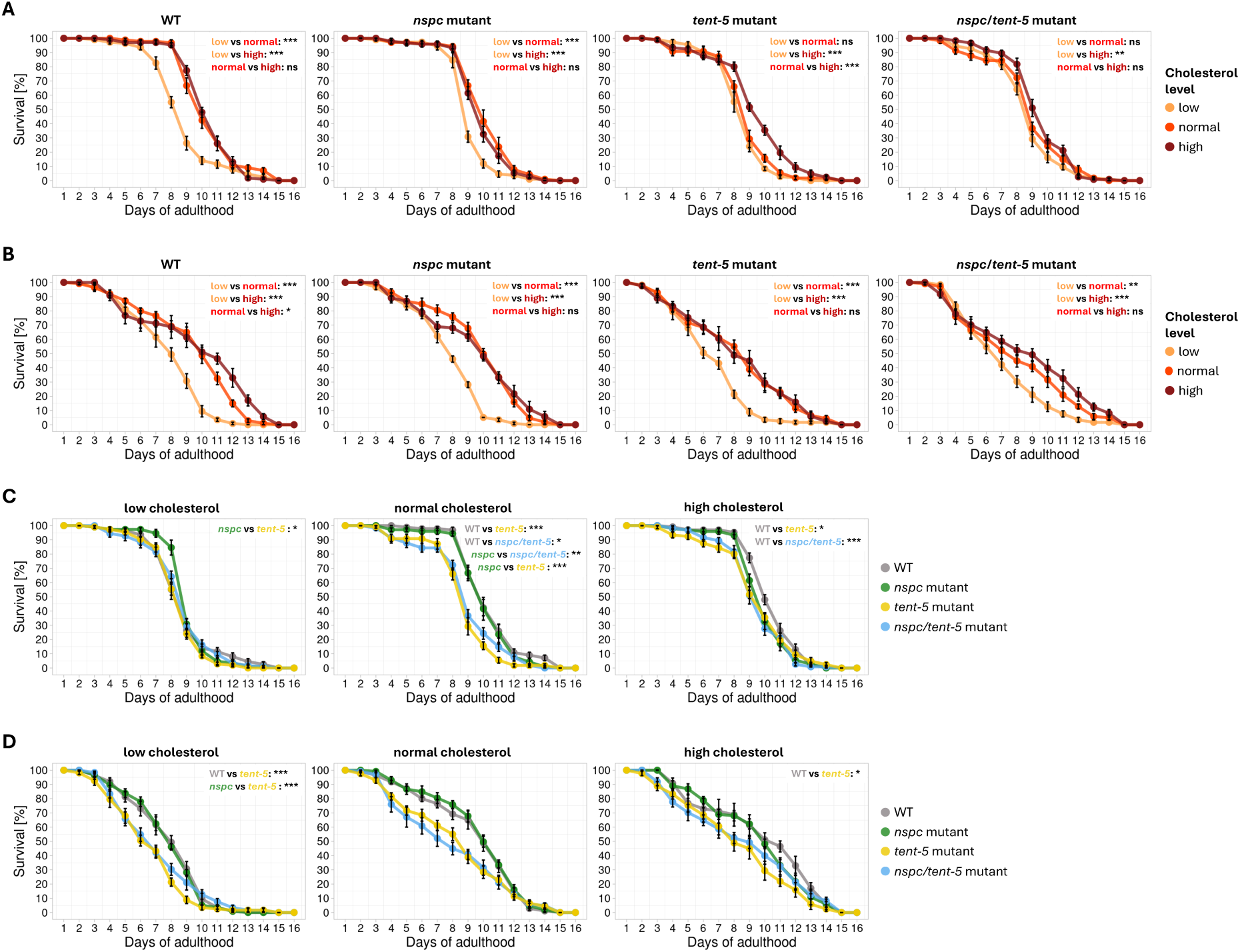
NSPCs might be involved in cholesterol metabolism. **(A)** Survival plots for wild-type, *nspc* mutant, *tent-5* mutant, and *nspc/tent-5* mutant worms grown on non-pathogenic *E. coli* HB101 plates supplemented with three different cholesterol concentrations: low (0 mg/ml; orange), normal (20 mg/ml; red), and high (80 mg/ml; brown). Points represent mean values from 6 separate plates (n≅20 each) with SD. ns => not significant; ** => *p*-value < 0.01; *** => *p*-value < 0.001 (log-rank test). **(B)** Survival plots for wild-type, *nspc* mutant, *tent-5* mutant, and *nspc/tent-5* mutant worms grown on pathogenic *P. aeruginosa* PAO1 plates supplemented with three different cholesterol concentrations: low (0 mg/ml; orange), normal (20 mg/ml; red), and high (80 mg/ml; brown). Points represent mean values from 6 separate plates (n≅20 each) with SD. ns => not significant; * => *p*-value < 0.05; ** => *p*-value < 0.01; *** => *p*-value < 0.001 (log-rank test). **(C)** Survival plots for low (0 mg/ml), normal (20 mg/ml), and high (80 mg/ml) cholesterol plates with non-pathogenic *E. coli* HB101 bacteria. Survival is compared among four strains: wild-type (gray), *nspc* mutant (green), *tent-5* mutant (yellow), and *nspc/tent-5* mutant (blue) worms. Points represent mean values from 6 separate plates (n≅20 each) with SD. Only significant comparisons are shown on the plots. * => *p*-value < 0.05; ** => *p*-value < 0.01; *** => *p*-value < 0.001 (log-rank test). **(D)** Survival plots for low (0 mg/ml), normal (20 mg/ml), and high (80 mg/ml) cholesterol plates with pathogenic *P. aeruginosa* PAO1 bacteria. Survival is compared among four strains: wild-type (gray), *nspc* mutant (green), *tent-5* mutant (yellow), and *nspc/tent-5* mutant (blue) worms. Points represent mean values from 6 separate plates (n≅20 each) with SD. Only significant comparisons are shown on the plots. * => *p*-value < 0.05; *** => *p*-value < 0.001 (log-rank test).

In conclusion, while the lifespan experiments are not straightforward, they do not rule out a potential role for TENT-5 and NSPCs in regulating cholesterol metabolism. This regulation could involve cholesterol uptake from the environment, its intracellular transport, or further chemical modifications.

### NSPC knockdown results in transcription signatures related to the DAF-2/DAF-16 insulin signaling

To further explore the potential functions of NSPCs, we analyzed high-throughput datasets from WormBase to identify conditions under which NSPCs become dysregulated (Supplementary Table S1). Our analysis showed that NSPCs are frequently dysregulated in studies involving the insulin DAF-2/DAF-16 pathway (Figure 6A). Following this lead, we examined the structures and sequences of NSPCs and noticed their similarity to insulin-like peptides from the INS family (Figure 6B and C). In *C. elegans,* these insulin peptides interact with the DAF-2 receptor, influencing the cytoplasmic or nuclear localization of the transcription factor DAF-16, which regulates key physiological processes such as lifespan, innate immunity, and metabolism [43–46] (Figure 6A). The similarity between NSPCs and insulin peptides led us to hypothesize that NSPCs might serve as neuropeptides that co-regulate the insulin signaling pathway, potentially in the excretory gland cell. Their potential redundancy with INS proteins could explain the lack of strong phenotypes following *nspc* cluster deletion. To test this hypothesis, we crossed a GFP-tagged DAF-16 strain [47] with both *tent-5* and *nspc* mutants and examined DAF-16-GFP localization. We observed no significant differences in DAF-16 localization across cell types in standard conditions, as well as after heat shock, pathogen infection, and varying cholesterol supply (Figure 6D, Supplementary Figure S5), suggesting that NSPCs and TENT-5 may not be essential for maintaining proper insulin signaling.

**Figure 6.**
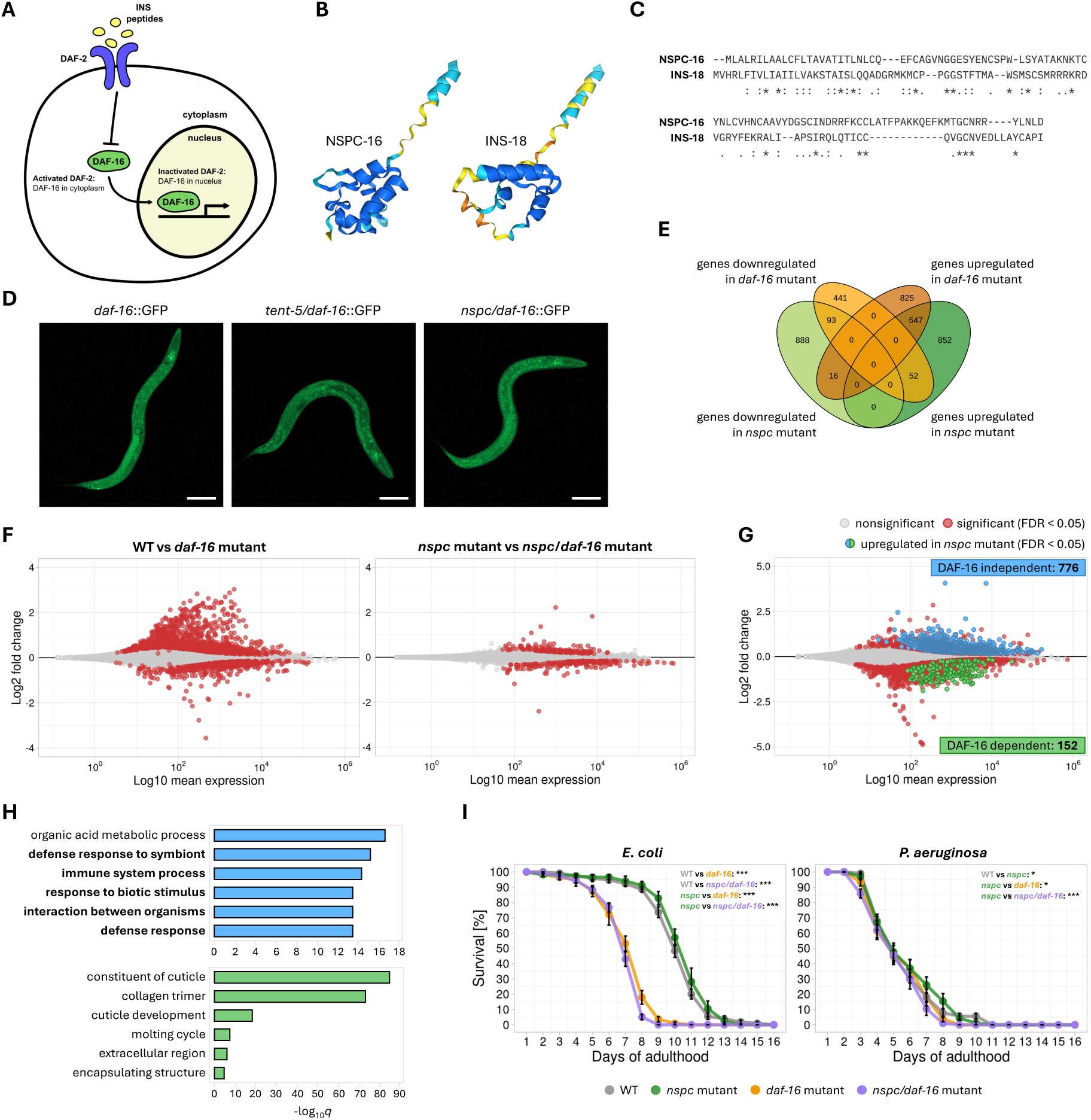
NSPCs might be involved in insulin signaling. **(A)** Schematic representation of the DAF-2/DAF-16 insulin signaling pathway. Insulin peptides (INS) act as ligands for the DAF-2 receptor. Upon DAF-2 activation, DAF-16 is retained in the cytoplasm; when DAF-2 is inactivated, DAF-16 translocates to the nucleus to initiate DAF-16-mediated gene expression. **(B)** Comparison of NSPC-16 and INS-18 protein structures predicted by AlphaFold 3 [48]. Representatives of NSPC and INS families were chosen arbitrarily. Protein similarity was assessed visually without the use of any quantitative parameters. **(C)** Comparison of NSPC-16 and INS-18 protein sequences. Asterisks (*) represent residues that are conserved in all NSPCs; colons (:) represent moderately conserved residues; and dots (.) indicate non-conserved residues with similar chemical properties. **(D)** Fluorescence microscopy images of DAF-16-GFP localization in wild-type, *tent-5*, and *nspc* mutant worms in standard growth conditions. Scale bars => 100 µm. **(E)** Venn diagram showing overlaps between upregulated (darker shade) and downregulated (lighter shade) genes in *daf-16* (orange) and *nspc* mutant (green) worms. Included are all significantly changed genes (FDR < 0.05). **(F)** MA plots showing differential gene expression after DAF-16 depletion in the presence of NSPCs (left) and in their absence (right). Significantly changed genes (FDR < 0.05) are marked with red dots. **(G)** MA plot showing differential gene expression between *daf-16* and *nspc/daf-16* mutant worms. Significantly changed genes (FDR < 0.05) are marked with red dots. Genes upregulated in *nspc* mutant that are DAF-16 independent are marked in blue, and DAF-16-dependent are shown in green. Only genes significantly upregulated in *nspc* mutant in both the presence (FDR < 0.05; Figure 4B; Supplementary Table S3) and absence of DAF-16 (Supplementary Table S5) were included. **(H)** Top GO terms for DAF-16 independent (top; blue) and dependent (bottom; green) genes upregulated in *nspc* mutant ordered by adjusted *p*-value. Terms related to the defense response are shown in bold. **(I)** Survival of wild-type (gray), *nspc* mutant (green), *daf-16* mutant (orange), and *nspc/daf-16* mutant (purple) worms grown on non-pathogenic *E. coli* HB101 (left) or pathogenic *P. aeruginosa* PAO1 (right). Points represent mean values from 6 separate plates (n≅20 each) with SD. Only significant comparisons are shown on the plots. * => *p*-value < 0.05; *** => *p*-value < 0.001 (log-rank test).

Next, we generated a strain lacking both *daf-16* and *nspc* genes and conducted RNA-seq to gain further insight into their possible co-regulation. Interestingly, we observed substantial overlap in genes upregulated upon deletion of either *nspc* genes or *daf-16* (Figure 6E, Supplementary Table S5), supporting the hypothesis of NSPC-mediated regulation of the insulin pathway and suggesting that NSPCs act antagonistically towards DAF-2. Consistently, our sequencing results showed that while DAF-16 depletion typically causes significant gene dysregulation, this effect is less pronounced in the absence of NSPCs (Figure 6F, Supplementary Table S5). In contrast, NSPC deletion in the absence of DAF-16 resulted in a transcriptomic shift similar to that seen with NSPC deletion alone, except for some genes that were upregulated when DAF-16 was present but downregulated in its absence (Figure 6G, Supplementary Table S5). These results suggest that NSPCs may act upstream of DAF-16, but their role is likely not limited to regulating insulin signaling. For example, defense response genes remained upregulated in the absence of NSPCs, regardless of DAF-16 presence (Figure 6H), indicating that these processes might be regulated independently of insulin signaling. Consequently, the deletion of NSPCs in *daf-16* mutants did not lead to any changes in worms’ survival for both non-pathogenic *E. coli* and pathogenic *P. aeruginosa* conditions (Figure 6I). Although our RNA-seq results support the potential role of NSPCs as neuropeptides in the DAF-2/DAF-16 pathway, their possible redundancy with other small peptides and the complexity of the signaling machinery might mask the potential physiological effect and make it difficult to definitively prove this hypothesis. A comprehensive study would require generating a strain lacking both NSPCs and all INS peptides; however, this is beyond the scope of the current work.

## Conclusions

The research presented in this manuscript revealed that one of the main functions of the excretory gland cell is the production of secreted nematode-specific NSPC proteins. This family, which encompasses 18 highly similar genes, is unique to the *Caenorhabditis* genus and was previously suggested to function as an immune effectors. Here, we show that they might function rather as neuropeptides that are potentially involved in regulating signaling pathways, such as the insulin DAF-2/DAF-16 pathway. Although these findings do not entirely clarify the physiological role of the excretory gland cell, they lay a foundation for future research into how this cell and its associated NSPCs contribute to nematode biology, particularly under ecologically relevant conditions.

From the evolutionary perspective, it is interesting to consider the function of the excretory gland cell in nematode species that lack NSPCs. Although limited studies exist, the excretory gland cell has been suggested to secrete components of epicuticle in *Toxocara canis* [49], enzymes important for feeding in *Nippostrongylus brasiliensi* [50], or peptidases required for molting in *Phocanema decipiens* [51]. While the first two possibilities remain unexplored in *C. elegans*, the role of the excretory gland cell in molting has already been excluded [11]. It appears that in *C. elegans*, the excretory gland cell has evolved to perform a distinct role, possibly because of inhabiting a different natural environment in comparison to the mentioned parasitic nematodes. That change was accompanied by the generation of *Caenorhabditis-* specific NSPCs, further supporting their importance for the cell’s function.

## Methods

### C. elegans culture and growth conditions

All *C. elegans* strains used in this study are summarized in Supplementary Table S6. Unless specified otherwise, worms were maintained at 20°C on nematode growth medium (NGM) plates seeded with *E. coli* HB101 as a food source.

### Plasmids construction

All plasmids used in this study are presented in Supplementary Table S7. Plasmids pVL033 and pVL037-40 were constructed using sequence- and ligation-independent cloning (SLIC), following established protocols. The VL033 plasmid was built by introducing *tent-5* in fusion with GFP-3xflag (amplified from respective BAC) into the pCFJ151 backbone. Plasmids pVL037-40 were assembled from the following fragments: the pCFJ104 backbone, the *nspc-9, −14, or −20* promoter with or without gene CDS (amplified from genomic DNA isolated from wild-type worms), the mCherry reporter (from pCFJ104), and the *nspc-9, −14, or −20* 3ʹ UTR (also from wild-type genomic DNA). PCR amplification of each fragment was performed using Phusion polymerase with an initial denaturation at 98°C for 2 min, followed by 25 cycles of 98°C for 10 s, 60°C for 30 s, and 72°C for 10 s, and a final elongation step at 72°C for 5 min. Plasmids pCE090 and pCE091 were prepared through classical restriction enzyme digestion and ligation methods. The *nspc-7* and *nspc-14* coding sequences were amplified from wild-type *C. elegans* cDNA and initially cloned into the pJET1.2 vector, following the manufacturer’s protocol. Both L4440 and pJET1.2 vectors containing *nspc-7* or *nspc-14* were digested with XhoI and XbaI and subsequently ligated with T4 DNA ligase. The pCE092 plasmid, containing *nspc-7*, *nspc-14*, and *nspc-20* sequences, was assembled with SLIC using an L4440 backbone containing the *nspc-14* sequence (from pCE091), *nspc-7* sequence (from pCE090), and *nspc-20* (from wild-type cDNA). All plasmids were validated by restriction digestion and Sanger sequencing.

### NSPC localization

Localization of NSPCs and TENT-5 was assessed using an Olympus FV1000 confocal microscope with a 60x/1.2 water immersion lens. Images were processed using Fiji/ImageJ software [52].

### Generation of transgenic C. elegans strains

All new *C. elegans* transgenic strains were generated by microinjection into the hermaphrodite gonad using either extrachromosomal arrays (plasmid-based) or CRISPR/Cas9-mediated genome editing (via guide RNA and repair template injection). Microinjections were conducted on Axio Observer 5 microscope.

Strains ADZ31, ADZ33, ADZ35, ADZ37, and ADZ44 were produced by injecting wild-type worms with a mix containing pVL037, pVL039, pVL038, pVL040, or pVL073 respectively (20 ng/μl), pRH269 (5 ng/μl), and GeneRuler DNA mix (75 ng/μl). Progeny were screened for green fluorescence in the pharynx and red fluorescence in the excretory gland cells, and transgenic worms were regularly enriched by isolating GFP-positive individuals. For the ADZ45 strain, pVL038 and pVL045 were injected without the co-injection marker.

Strains ADZ72, ADZ83, and ADZ111-114 were generated using a modified co-CRISPR approach, based on established protocols. Injection mixes contained 4 μM Cas9, 10 μM specific guide/s, 4 μM *dpy-10* guide RNA, 0.4 μM repair template, and 0.2 μM *dpy-10* repair template. To create strain ADZ72, the *nspc-10* coding sequence was removed using two guide RNAs and repaired by a template containing the first *nspc-10* intron, *tomm-20*, and miniSOG(426Cys) sequences. The ADZ82 strain was generated by introducing a single nucleotide substitution in miniSOG present in ADZ72, converting Cys426 to Gly and thus activating the miniSOG protein. For the ADZ111 strain, the first NSPC cluster was removed, and successive deletions were conducted until the final strain lacking all NSPC clusters was obtained. In all cases, worms were co-injected with *dpy-10* guide and repair template introducing a visible roller or dumpy phenotype in affected worms, enabling easier selection of mutants for genotyping. After homozygote selection, worms were outcrossed with wild-type males to eliminate any potential off-target mutations. All CRISPR guide RNAs and repair templates are listed in Supplementary Table S8. The repair template for strain ADZ72 was generated by PCR from pCZGY1703 plasmid, using primers with 35-50 nucleotide homology arms. The PCR reaction was performed using Phusion polymerase with an initial denaturation at 98°C for 2 min, followed by 25 cycles of 98°C for 10 s, 60°C for 30 s, and 72°C for 10 s, and a final elongation step at 72°C for 5 min. PCR products were purified using Gel-out and KAPA Pure magnetic beads, and prepared for microinjection.

Additional strains, including ADZ74, ADZ83, ADZ116, ADZ122, ADZ130, and ADZ131, were generated by crossing previously established strains as outlined in Supplementary Table S6.

### Optogenetic ablation

To induce optogenetic cell ablation, age-synchronized populations of wild-type and active/inactive miniSOG worms were cultured on NGM plates at 15°C until the L3 stage. Plates were then exposed to blue light (460 nm) from an LED advertising panel. The plates were inverted and placed without lids on the panel, with a 1 mm glass separator to minimize the transfer of heat. The illumination process was performed at ambient temperatures of 18-20°C and included five intervals of 10 minutes of light, separated by 10-minute breaks. After illumination, worms were transferred back to 15°C and grown for two days to allow complete miniSOG-mediated cell destruction. Ablation success was verified by detecting the loss of mCherry fluorescence in the excretory gland cell and measuring the expression of *nspc* genes by RT-qPCR. All imaging was conducted on a Zeiss LSM800 confocal microscope with a 40×/oil immersion lens.

### Western blotting

Similarly dense population of worms expressing active or inactive miniSOG were harvested and washed three times with 50 mM NaCl. Worm pellets were mixed with zirconia-silica beads and 3x SDS sample buffer (187.5 mM Tris-HCl (pH 6.8), 6% SDS, 150 mM DTT, 0.02% bromophenol blue, 30% glycerol, and 3% 2-mercaptoethanol), vortexed for 5 minutes, and incubated at 95°C for 5 minutes. Samples were centrifuged, and the supernatant was run on 15% SDS-PAGE gels. Proteins were transferred onto Protran nitrocellulose membranes through semi-dry transfer at 15V for 1 hour. Following the transfer, membranes were blocked with 5% non-fat milk in 1x TBST (20 mM Tris-HCl, 150 mM NaCl, 0.01% Tween 20) for 1 hour at room temperature. Membranes were then incubated with primary antibodies for miniSOG (1:1000; Kerafast; EFH004) and α-tubulin (1:10 000; Sigma-Aldrich; CP06) overnight at 4°C. After washing with 1x TBST, membranes were incubated with secondary anti-mouse antibodies (1:5000 for miniSOG, 1:10000 for α-tubulin; Sigma-Aldrich; 401215) for 1 hour at room temperature, activated with Clarity Western ECL Substrate (Bio-Rad), and visualized using CL-Exposure film (Thermo Fisher Scientific) developed with an AGFA Curix CP-1000 device.

### RNA interference (RNAi)

RNA interference (RNAi) was used to silence NSPC genes in *C. elegans*. Plates for RNAi were additionally supplemented with 1 mM IPTG (Sigma-Aldrich), 25 μg/ml carbenicillin (Sigma-Aldrich), and 25 U/ml nystatin (Sigma-Aldrich). *E. coli* HT115 bacteria transformed with RNAi plasmids were prepared by overnight culture, centrifuged, concentrated 15x times, and seeded onto RNAi plates, which were left at room temperature overnight. Age-synchronized L4 worms (approximately 30 per 100-mm plate) were transferred to the RNAi plates, and their progeny were used for subsequent analyses. Silencing efficiency was verified using RT-qPCR.

### RNA isolation

Populations of worms were collected and washed three times with 50 mM NaCl. Worm pellets were resuspended in 1 ml TRI Reagent (Sigma-Aldrich), vortexed for 15 minutes at room temperature, and stored at −80°C until further processing. RNA extraction was performed according to the manufacturer’s protocol for TRIzol-based isolation, with additional purification using KAPA Pure magnetic beads (RNA to beads ratio: 1:3 v/v). RNA quality and integrity were confirmed using the Agilent TapeStation system. RNA samples were prepared in three independent biological replicates for downstream RT-qPCR and Illumina RNA-seq, and two replicates were used for direct RNA sequencing.

### RT-qPCR

Expression levels of NSPCs following excretory gland cell ablation and RNAi silencing were evaluated using RT-qPCR. Total RNA was treated with TURBO DNase (Thermo Fisher Scientific) to eliminate genomic DNA. cDNA synthesis was conducted following the manufacturer’s protocol with the use of SuperScript III (Thermo Fisher Scientific) and a mix of oligo(dT)_20_ and random primers. RT-qPCR reactions were carried out using Platinum SYBR Green Mix (Thermo Fisher Scientific) on a QuantStudio 5 PCR system (Thermo Fisher Scientific) with primers targeting NSPCs (Supplementary Table S8). Due to high similarity in NSPC sequences, primers were designed to detect groups of genes: one primer pair for *nspc- (1-15),* and another for *nspc-(16-20).* Relative gene expression was normalized to *act-1* or *rps-23* and quantified using the 2^−ΔΔC(t)^ method. Statistical significance was determined by two-tailed unpaired Student’s t-tests.

### Illumina RNA sequencing and data analysis

RNA samples were treated with TURBO DNase (Thermo Fisher Scientific) before sequencing. Libraries were prepared from 2 μg of poly(A)-enriched RNA and sequenced on an Illumina platform in 150-bp paired-end mode (performed by the Center for New Technology, University of Warsaw or GENEWIZ, Germany). Reads were mapped to the *C. elegans* WBCel235 reference genome using STAR aligner v2.7.10a [53] and processed with samtools v1.961 [54]. Read counts were obtained with featureCounts from the Subread package v2.0.6 [55] (options -Q 10 -p -C -B). Differential expression analysis was conducted using DESeq2 Bioconductor package v1.28 [56] with default settings. Gene ontology (GO) enrichment analysis was performed using WormBase Enrichment Suite [57]. Raw sequencing data were deposited in the Gene Expression Omnibus (GEO) database under accession number GSE282286.

### Nanopore Direct RNA sequencing and data analysis

Libraries were generated using the Direct RNA Sequencing Kit (SQK-RNA002, Oxford Nanopore Technologies), using 1.5 μg of total RNA with added in vitro transcribed poly(A) standards (2 ng). Sequencing was conducted on a MinION device and followed by basecalling by Guppy v6.0.0. Reads were aligned to the WBCel235 reference using MiniMap v2.17 [58] (options -k 14 -ax map-ont –secondary=no) and processed with samtools v1.9 [54]. Poly(A) tail lengths were estimated using Nanopolish v0.13.2 as described before [13,59]. Distribution comparisons across conditions were assessed by the Wilcoxon test, with p-values adjusted via the Benjamini-Hochberg method. Raw DRS data are available in the GEO database under accession number GSE282286.

### Worms age estimations

Worm age was estimated for samples used in RNA-seq experiments using the R packages RAPToR v1.2 and wormRef v0.5 [26]. Given the worms’ stages (L4 for NSPC mutants and young adults for optogenetic ablations), the *“Cel_YA_1”* reference was selected from wormRef as appropriate for both experiments. Age estimations were conducted using *ae* function with normalized gene counts as an input. Statistical differences were tested by a two-tailed unpaired Student’s t-test.

### Lifespan experiments

Lifespan assays were conducted on 35-mm NGM plates supplemented with 50 μM 5-fluoro-2′-deoxyuridine (FUdR) to inhibit reproduction and maintained at 25°C. To assess NSPC function in cholesterol metabolism, plates were prepared with varying cholesterol concentrations of 0 mg/ml (low), 20 mg/ml (standard), and 80 mg/ml (high). All plates were seeded with either non-pathogenic *E. coli* HB101 or pathogenic *P. aeruginosa* PAO1 (100 μl/plate) and incubated overnight at 37°C. Age-synchronized *C. elegans* populations were prepared by bleaching and seeding embryos on plates. Once worms reached the L4 stage, they were transferred to the experimental plates (20 worms per plate, six replicates per condition). The number of live and dead worms was recorded daily until all worms had died. Survival data were analyzed by the Kaplan-Meier method, with condition comparisons assessed via log-rank significance tests.

### Brood-size analysis

To evaluate reproductive potential post-ablation, progeny counts were done for wild-type, inactive miniSOG (ADZ74), and active miniSOG (ADZ83) worms, with and without illumination. Worms were synchronized by bleaching, grown to L3 at 15°C, and half of the population was illuminated as described. After 24 hours, five worms per condition were placed on fresh plates seeded with *E. coli* HB101 and later grown at 20°C. Worms were transferred to new plates daily throughout their entire lifespan. Progeny counts were compared across conditions by a two-tailed unpaired Student’s t-test. For RNAi-silenced worms, assays followed a similar approach but with HT115 RNAi bacterial plates and worms’ preparation steps used in RNAi protocol.

### DAF-16 localization

Localization of DAF-16 was visualized in strains OH16024 (*daf-16*::*GFP)* [47], ADZ116 (*tent-5/daf-16::GFP*), and ADZ122 (*nspc/daf-16::GFP*) under standard conditions upon various stimuli. For standard imaging, synchronized L4 worms were immobilized with 25 μM levamisole on freshly prepared 2% agarose pads and imaged immediately. For studying DAF-16 nuclear translocation after heat shock, synchronized L4 worms were exposed to 35°C for 1 hour before imaging. For studying DAF-16 delocalization from the nucleus to the cytoplasm upon infection, worms after heat shock were either maintained on standard *E. coli* HB101 or switched to *P. aeruginosa* PAO1 for 16 hours before imaging as described before [60]. To investigate the effects of cholesterol concentration on DAF-16 localization, worms were grown on NGM plates containing low (0 mg/ml) or high (80 mg/ml) cholesterol before standard imaging. All imaging was conducted on a Zeiss LSM800 confocal microscope with a 20x objective.

### Sequence alignment and phylogenetic analysis

Amino acid sequences of all NPSC and INS peptides were retrieved from WormBase [19]. Alignments and phylogenetic analysis were performed using the MUSCLE tool from EMBL-EBI [20].

## Data availability

The raw data from Illumina RNA Sequencing and Nanopore Direct RNA Sequencing have been deposited in the Gene Expression Omnibus (GEO) database under the accession number GSE282286.

## Acknowledgments

We are grateful to the members of the Dziembowski group and Krzysztof Drabikowski for the discussion and valuable feedback. We especially thank Paweł Krawczyk for helping with bioinformatic analyses, Aleksandra Brouze and Seweryn Mroczek for running Nanopore DRS sequencing, Jonathan J. Ewbank and Krzysztof Drabikowski for sharing plasmids, and Tomasz Węgierski for supporting microscopic observations.

## Author contributions

A.D. provided funding and directed the project. Z.M., V.L., and A.D. designed the experiments. V.L. generated the strains ADZ31-ADZ45 and performed visualizations of NSPC localization and its colocalization with TENT-5. Z.M. prepared the remaining strains, carried out the remaining experiments, and analyzed all sequencing data. Z.M. prepared the initial draft of the manuscript and created all figures. V.L. and A.D. revised and edited the manuscript.

