## Supplementary Materials for "Knockout of all nematode-specific NSPC genes expressed exclusively in the excretory gland cell results in transcriptomic signatures indicating an affected insulin signaling"

(Supplementary Figures S1-S5, Supplementary Tables S6-S8)

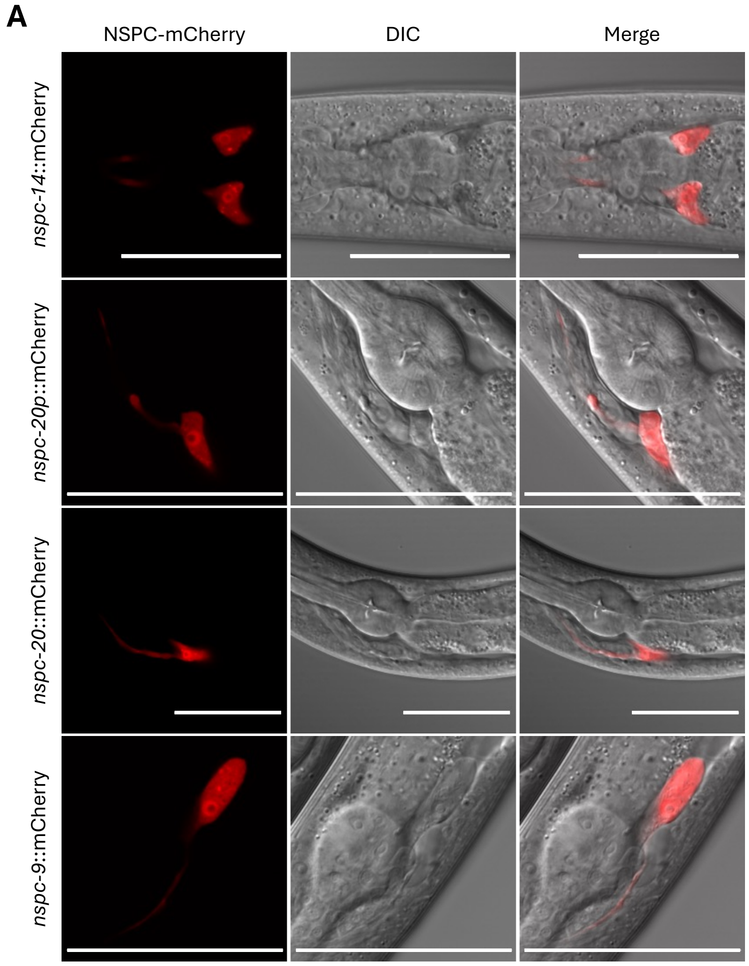


**Supplementary Figure S1. NSPCs are exclusively expressed in the excretory gland cell.**
**(A)** Additional fluorescence and differential interference contrast (DIC) microscopy images of NSPC localization in the excretory gland cell for different transcriptional and translational NSPC-mCherry reporters. Scale bars => 50 µm.

**A**

| Blue light source | Mode of light | Worms' stage | Illumination time | Temperature | Ablation efficiency | Additional comments |
|---|---|---|---|---|---|---|
| stereo microscope fluorescence adapter (440-460 nm) | continuous light | multiple variants | multiple variants | 20°C | depending on condition | - illumination of one plate at a time<br>- uneven illumination of plate |
| military torches (460 nm and 470 nm) | continuous light | | | | | |
| WormLab's optogenetic module (470 nm) | flushing light | | | | | |
| LED board (460 nm) | continuous light; flushing light | L4 | 30 min;<br>10 min | 20°C | medium | - better effect for continuous light<br>- better effect for longer illumination |
| LED board (460 nm) | continuous light | L4 | 1 h | 20°C | medium | - many dead worms |
| LED board (460 nm) | continuous light | L4 | 45 min | 15°C | low | |
| LED board (460 nm) | continuous light | L2 | 30 min | 20°C | low | |
| LED board (460 nm) | continuous light | embryo-L4 | 5 min every 2 h | RT | low | - worms' developmental delay |
| LED board (460 nm) | continuous light | L4 | 30 min (repeated next day);<br>2x15 min (repeated next day) | 20°C | high | |
| **LED board (460 nm)** | **continuous light** | **L3** | **5x10 min** | **15°C** | **high** | **- complete disappearance of the excretory gland cell** |

**B**

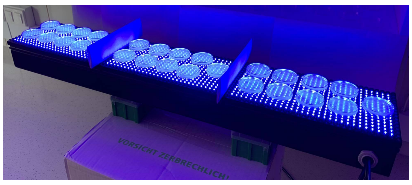

**C**

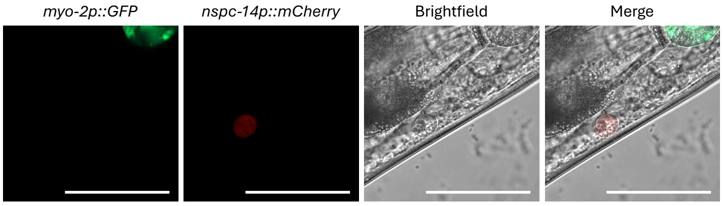


**Supplementary Figure S2. Optogenetic ablation can be implemented to destroy the excretory gland cell.** **(A)** Table summarizing the optimization process for optogenetic ablation of the excretory gland cell. Various blue light sources, illumination durations, modes of light, and developmental stages of the worms were tested.
**(B)** LED advertising board used for illumination of miniSOG-expressing worms. The LED board emits 460 nm blue light with adjustable frequency, allowing simultaneous illumination of multiple plates containing worms.
**(C)** Fluorescence microscopy image of the excretory gland cell after optogenetic ablation. In some cases, the excretory gland cell was not destroyed completely but was significantly shrunken. However, such a drastic change in cell size and morphology is likely sufficient to impair its functions. Scale bars => 50 µm.

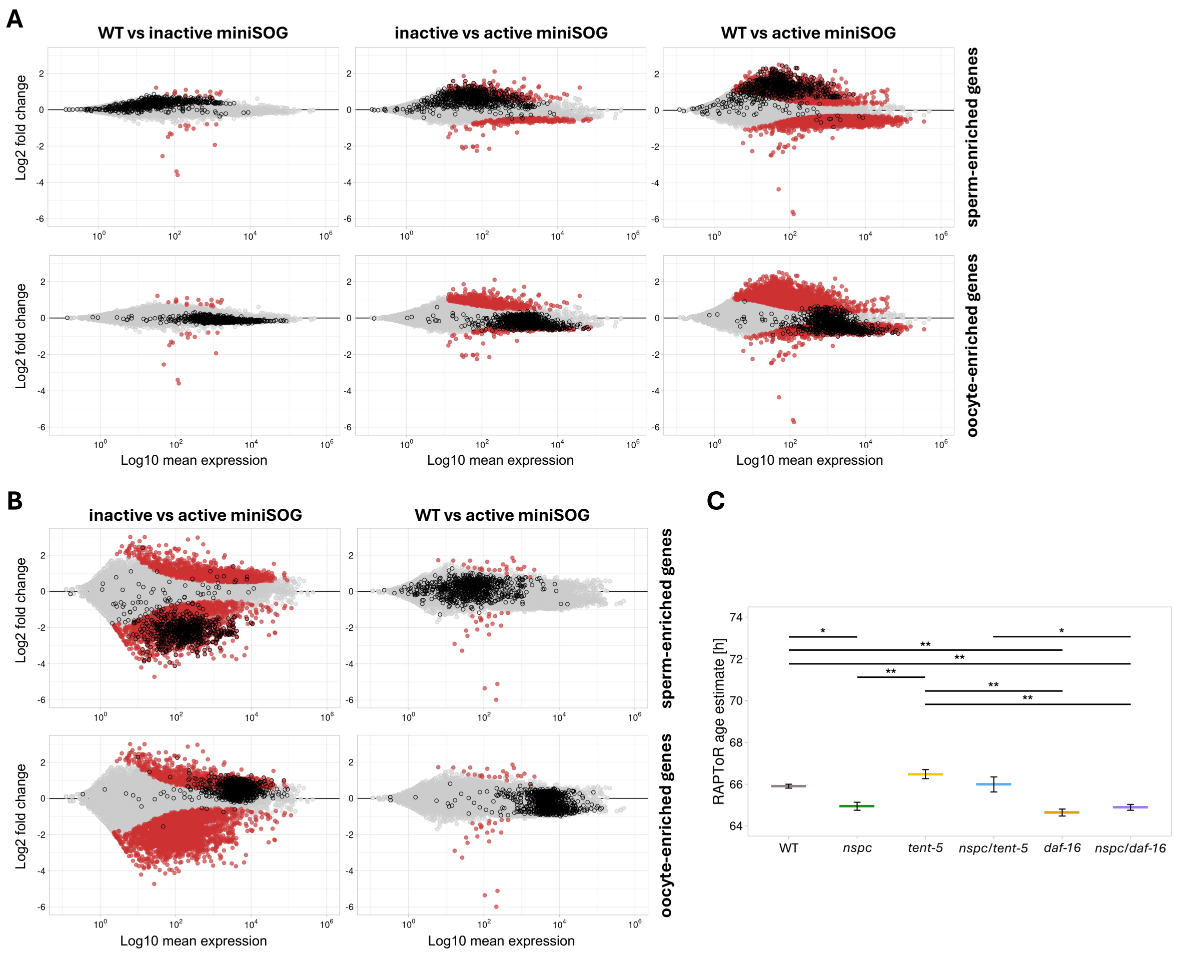


**Supplementary Figure S3. Ablation of the excretory gland cell leads to no transcriptome changes.**

**(A)** MA plots illustrating all possible combinations of differential gene expression between wild-type, inactive, and active miniSOG worms from the second replicate of the excretory gland cell ablation. Significantly changed genes (FDR < 0.05) are marked with red dots. Black borderline is used to mark sperm- and oocyte-enriched genes, as in Figure 3C.

**(B)** MA plots illustrating all remaining combinations of differential gene expression between wild-type, inactive, and active miniSOG worms from the third replicate of the excretory gland cell ablation. Significantly changed genes (FDR < 0.05) are marked with red dots. Black borderline is used to mark sperm- and oocyte-enriched genes, as in Figure 3C.

**(C)** Age estimates for wild-type (gray), *nspc* mutant (green), *tent-5* mutant (yellow), *nspc/tent-5* mutant (blue), *daf-16* mutant (orange), and *nspc/daf-16* mutant (purple) worms from other RNA-seq experiments described in this study. Although age differences are significant between multiple conditions, they are much smaller than those observed in RNA-seq samples after the ablation. Plots represent mean values with SD. Only significant comparisons are shown on the plots. ns => not significant; * => *p*-value < 0.05; ** => *p*-value < 0.01 (two-tailed *t*-test).

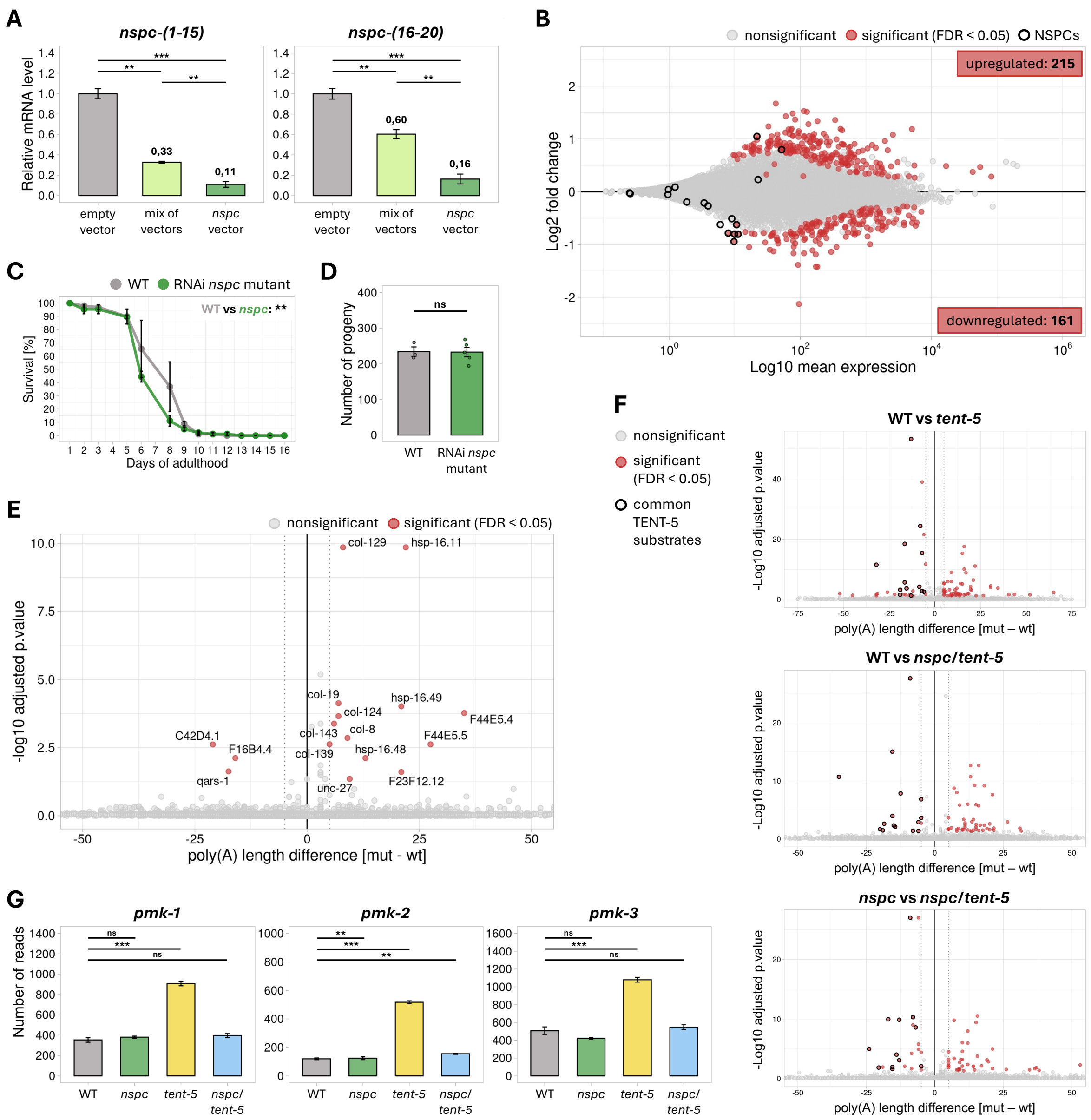


**Supplementary Figure S4. NSPC deletion does not result in any physiological phenotypes.**

**(A)** RT-qPCR showing decreased expression levels of all *nspc* genes following RNAi silencing. Worms were grown on *E. coli* HT115 expressing one of the following: an empty RNAi vector (control; gray), a mix of bacteria each expressing a vector targeting *nspc-7*, *-14*, or *-20* individually (light green), or a single silencing vector simultaneously targeting *nspc-7, -14,* and *-20* (*nspc-7/14/20*) (green). Relative *nspc* mRNA levels were normalized to *act-1*. Bar plots represent mean values with SD. ** => p-value < 0.01; *** => p-value < 0.001 (two-tailed t-test). Due to higher efficiency, the *nscp-7/14/20* vector was used for further RNAi silencing experiments.

**(B)** MA plot illustrating differential gene expression between wild-type and RNAi *nspc* mutant worms grown on *E. coli* HT115 expressing empty RNAi vector or *nspc-7/14/20* silencing vector, respectively. Significantly changed genes (FDR < 0.05) are marked with red dots (Supplementary Table S3). Black borderline is used to mark *nspc* genes. *Nspc-7*, *-14*, and *-20* were upregulated despite RNAi silencing due to the detection of bacterial dsRNA during RNA-seq.

**(C)** Survival of wild-type (gray) and RNAi *nspc* mutant (green) worms grown on *E. coli* HT115 expressing empty RNAi vector or *nspc-7/14/20* silencing vector, respectively. Points represent mean values from 3 separate plates (n≅40 each) with SD. ** => *p*-value < 0.01 (log-rank test).

**(D)** Differences in brood sizes between wild-type (gray) and RNAi *nspc* mutant (green) worms grown on *E. coli* HT115 expressing empty RNAi vector or the *nspc-7/14/20* silencing vector, respectively. Bar plots represent mean values with SD (n=3 for WT, n=5 for RNAi *nspc* mutant). ns => not significant (two-tailed t-test).

**(E)** Volcano plot showing differential polyadenylation between wild-type and *nspc* mutant worms (Supplementary Table S4). Transcripts with significantly changed median poly(A) tail length (FDR < 0.05) by a minimum of 5 nucleotides (dotted lines) are marked with red dots and labeled.
**(F)** Volcano plots showing a lack of NSPC influence on differential polyadenylation between wild-type and *tent-5* mutant worms (Supplementary Table S4). Transcripts with significantly changed median poly(A) tail length (FDR < 0.05) by a minimum of 5 nucleotides (dotted lines) are marked with red dots. Black borderline is added to transcripts commonly shortened in all three comparisons.
**(G)** Plots showing the number of RNA-seq reads for *pmk* genes in wild-type (gray), *nspc* mutant (green), *tent-5* mutant (yellow), and *nspc/tent-5* mutant (blue) worms. Bar plots represent mean values with SD. ns => not significant; ** => p-value < 0.01; *** => p-value < 0.001 (two-tailed t-test).

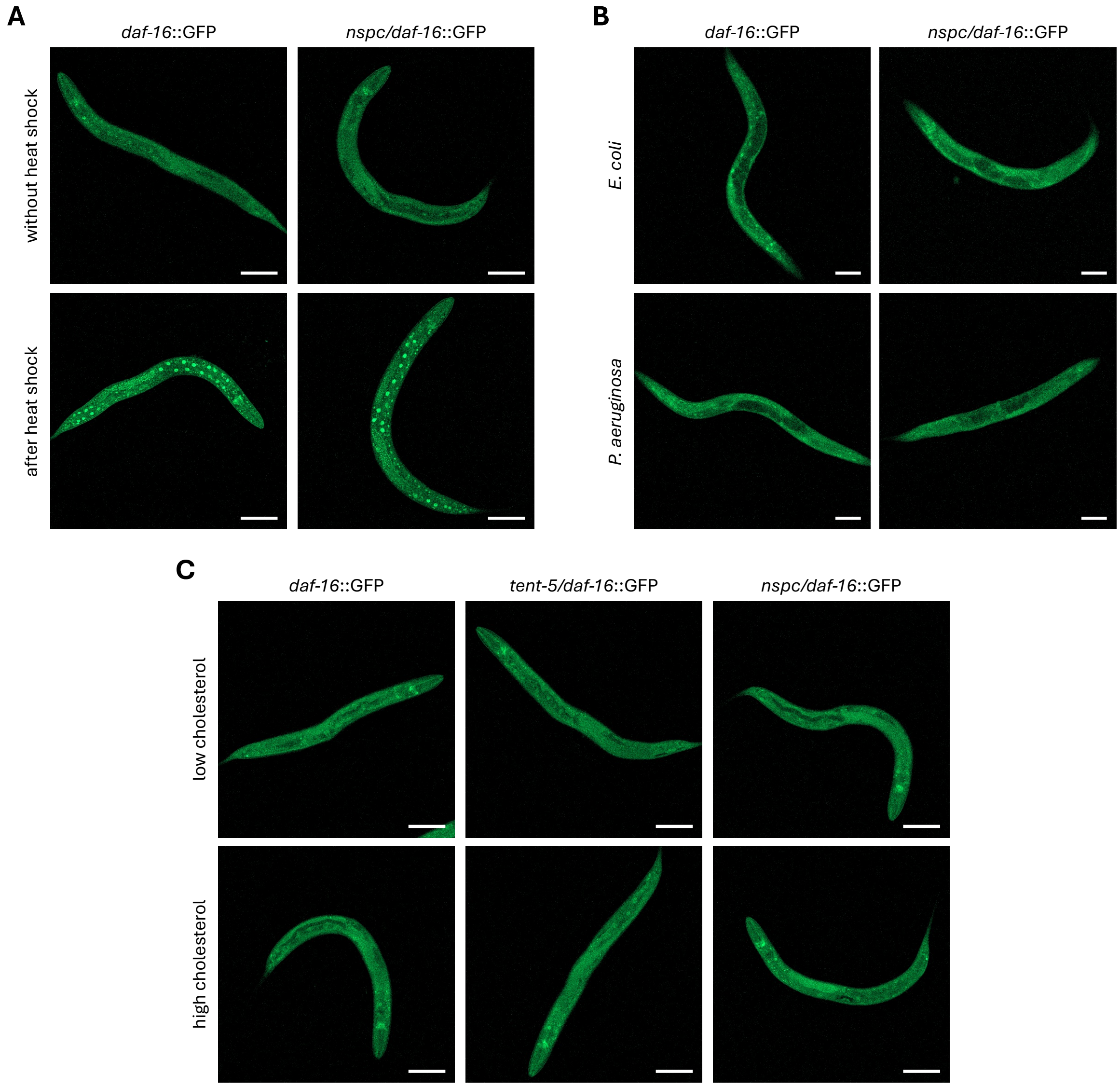


**Supplementary Figure S5. Studying the influence of TENT-5 and NSPCs on DAF-16 localization.**
**(A)** Fluorescence microscopy images of DAF-16-GFP localization in wild-type and *nspc* mutant worms after heat shock (1h; 35ºC), which is the best-known stimuli for relocating DAF-16 from the cytoplasm to the nucleus. Scale bars => 100 µm.
**(B)** Fluorescence microscopy images of DAF-16-GFP localization in wild-type and *nspc* mutant worms after heat shock (1h; 35ºC) followed by exposure to either non-pathogenic *E. coli* HB101 or pathogenic *P. aeruginosa* PAO1 as described in Evans E.A. *et al.* (2008) [60]. Scale bars => 100 µm.
**(C)** Fluorescence microscopy images of DAF-16-GFP localization in wild-type, *tent-5*, and *nspc* mutant worms grown on plates with low (0 mg/ml) or high (80 mg/ml) cholesterol concentration. Scale bars => 100 µm.

**Supplementary Table S6. List of *C. elegans* strains used in this study.**

| Strain | Genotype | Source | Comments |
|---|---|---|---|
| N2 Bristol | wild type | CGC* | - |
| CF1038 | *daf-16(mu86) I* | CGC* | - |
| OH16024 | *daf-16(ot971[daf-16::GFP]) I* | CGC* | - |
| TM3504 | *tent-5(tm3504) I* | NBRP** | - |
| ADZ31 | *rttEx12[nspc-14p::mCherry::nspc-14 3' UTR; myo-2p::gfp::unc-54 3' UTR] in N2* | this study | extrachromosomal array; transcriptional reporter |
| ADZ33 | *rttEx13[nspc-20p::mCherry::nspc-20 3' UTR; myo-2p::gfp::unc-54 3' UTR] in N2* | this study | extrachromosomal array; transcriptional reporter |
| ADZ35 | *rttEx15[nspc-14p::nspc-14::mCherry::nspc-14 3' UTR; myo-2p::gfp::unc-54 3' UTR] in N2* | this study | extrachromosomal array; translational reporter |
| ADZ37 | *rttEx17[nspc-20p::nspc-20::mCherry::nspc-20 3' UTR; myo-2p::gfp::unc-54 3' UTR] in N2* | this study | extrachromosomal array; translational reporter |
| ADZ44 | *rttEx24[nspc-9p::nspc-9::mCherry::nspc-9 3' UTR; myo-2p::gfp::unc-54 3' UTR] in N2* | this study | extrachromosomal array; translational reporter |
| ADZ45 | *rttEx25[tent-5p::tent-5abd::egfp::3xflag::tent-5 3' UTR; nspc-14p::nspc-14::mCherry::nspc-14 3' UTR] in N2* | this study | extrachromosomal array |
| ADZ72 | *nspc-10(rtt31[nspc-10p::tomm-20::miniSOG(426Cys)::nspc-10 3' UTR]) X* | this study | CRISPR/Cas9 |
| ADZ74 | *rttEx12[nspc-14p::mCherry::nspc-14 3' UTR; myo-2p::gfp::unc-54 3' UTR] in ADZ72* | this study | cross between ADZ72 and ADZ31; **final inactive miniSOG strain** |
| ADZ82 | *nspc-10(rtt33[nspc-10p::tomm-20::miniSOG::nspc-10 3' UTR]) X* | this study | CRISPR/Cas9 |
| ADZ83 | *rttEx28[nspc-14p::mCherry::nspc-14 3' UTR; myo-2p::gfp::unc-54 3' UTR] in ADZ82* | this study | cross between ADZ82 and ADZ31; **final active miniSOG strain** |
| ADZ111 | *nspc-(1-7)(rtt34) X* | this study | CRISPR/Cas9 |
| ADZ112 | *nspc-(1-10)(rtt35) X* | this study | CRISPR/Cas9 |
| ADZ113 | *nspc-(1-15)(rtt36) X* | this study | CRISPR/Cas9 |
| ADZ114 | *nspc-(1-20)(rtt37) X* | this study | CRISPR/Cas9; **final *nspc* mutant** |
| ADZ116 | *tent-5(tm3504) I; daf-16(ot971[daf-16::GFP]) I* | this study | cross between TM3504 and OH16024 |
| ADZ122 | *nspc-(1-20)(rtt37) X; daf-16(ot971[daf-16::GFP]) I* | this study | cross between ADZ114 and OH16024 |
| ADZ130 | *nspc-(1-20)(rtt37) X; tent-5(tm3504) I* | this study | cross between ADZ114 and TM3504 |
| ADZ131 | *nspc-(1-20)(rtt37) X; daf-16(mu86) I* | this study | cross between ADZ114 and CF1038 |

***CGC** - Caenorhabditis Genetics Center
****NBPJ** - National Bioresource Project of Japan

**Supplementary Table S7. List of plasmids used in this study.**

| ID | Sequence | Source |
|---|---|---|
| pCFJ104 | *myo-3p::mCherry:unc-54 3' UTR* | J. J. Ewbank |
| pRH269 | *myo2p::gfp::unc-54 3' UTR* | K. Drabikowski |
| pCZGY1703 | *tomm-20N::miniSOG(426Cys)* in pENTR | Addgene #66781 |
| L4440 | - | K. Drabikowski |
| pCFJ151 | - | J. J. Ewbank |
| pJET1.2 | - | Thermo Fisher Scientific |
| WRM069A | BAC containing *tent-5abcd::egfp::3xflag* | TransgeneOme |
| pVL033 | *tent-5pshort::tent-5abd::egfp::3xflag::tent-5 3' UTR* in pCFJ151 | this study |
| pVL037 | *nspc-14p::mCherry::nspc-14 3' UTR* in pCFJ104 | this study |
| pVL038 | *nspc-14p::nspc-14::mCherry::nspc-14 3' UTR* in pCFJ104 | this study |
| pVL039 | *nspc-20p::mCherry::nspc-20 3' UTR* in pCFJ104 | this study |
| pVL040 | *nspc-20p::nspc-20::mCherry::nspc-20 3'UTR* in pCFJ104 | this study |
| pVL073 | *nspc-9p::nspc-9::mCherry::nspc-9 3' UTR* in pCFJ104 | this study |
| pCE090 | *nspc-7* in L4440 | this study |
| pCE091 | *nspc-14* in L4440 | this study |
| pCE092 | *nspc-7::nspc-14::nspc-20* in L4440 | this study |

**Supplementary Table S8. List of guide RNAs, repair templates and primers used in this study.**

| ID | Sequence | Comments |
|---|---|---|
| ***guide RNAs*** | | |
| sgRNA01 | GCUACCAUAGGCACCACGAG | *dpy-10*; for co-CRISPR |
| sgRNA08 | AAAGGACAACGAGGGTGCGA | for ADZ72 generation |
| sgRNA09 | AGCTTTACAACAATCTTTAA | |
| sgRNA17 | TCTCTGGTCCCTGCAGAAAG | for ADZ82 generation |
| sgRNA18 | TATAGATGTGATATAATGAC | for ADZ111 generation |
| sgRNA19 | AGGAAGTGGATGTTATCTCC | |
| sgRNA20 | ATTGTTCTAGATGCGCAAAC | for ADZ112 generation |
| sgRNA21 | ACAGCTCCTGTTATCAATAC | |
| sgRNA22 | TTGTGATCGCATTTTAAATG | for ADZ113 generation |
| sgRNA23 | CATCACTTGAACAGTAATCC | |
| sgRNA24 | TGTAACCAGGCCGTGTTTGA | for ADZ114 generation |
| sgRNA25 | GTGAATACCACCCACCAGTT | |
| ***repair templates*** | | |
| VL314 | CACTTGAACTTCAATACGGCAAGATGAGAATGACTGGAA<br>ACCGTACCGCATGCGGTGCCTATGGTAGCGGAGCTT<br>CACATGGCTTCAGACCAACAGCCTAT | *dpy-10*; for co-CRISPR |
| - | C**ATG**GTAAGTAGTTTCAGTTTTAAATGGAACAATTGAAT<br>ATCCTTGCAGTCGGACACAATTCTTGGTTTCAACAAATCA<br>AACGTCGTTTTGGCTGCTGGAATTGCTGGAGCCGCTTTCC<br>TCGGCTACTGCATTTACTTCGATCATAAGAGAATCAACGC<br>TCCAGACTACAAGGACAAGATTAGGCAAAAGAGACGTGC<br>CCAGGCTGGAGCATCCGGAGAGAAAAGTTTCGTGATAAC<br>TGATCCACGGCTGCCAGACAATCCCATCATCTTCGCATCC<br>GATGGCTTCCTGGAGCTGACCGAGTATTCCAGAGAGGAG<br>ATCCTGGGCCGCAAT**T**GCCGCTTTCTGCAGGGACCAGAG<br>ACAGACCAGGCCACAGTGCAGAAGATTCGCGATGCCATT<br>AGAGATCAGCGCGAGATTACCGTGCAGCTGATAAACTAC<br>ACAAAAAGCGGGAAGAAATTCTGGAACCTCCTGCACCTC<br>CAGCCCATGAGGGACCAGAAGGGTGAGCTCCAGTATTTC<br>ATCGGAGTGCAGCTGGATGGAGGATCTTCCGGATAAAGG<br>AGTTTTCTGCACATTCAGTTGTTGCACATTCATTC | for ADZ72 generation<br><br>**ATG** – start codon<br>XXX – *nspc-10* intron<br>XXX – *tomm-20*<br>XXX – *miniSOG(426Cys)*<br>**T** – place of point mutation (G in active *miniSOG*) |
| prCE580 | GTATTCCAGAGAGGAGATCCTGGGCCGCAATGGTCGCTT<br>TCTGCAGGGACCAGAGACAGACC | for ADZ82 generation |
| prCE660 | TAAACCAGATTTGTAAATTATAGATGTGATATAATTCCTG<br>GTTTGCTGAAAATTATTTGCAATATGTGCC | for ADZ111 generation |
| prCE661 | TCCCGCAAACAAAAATACATTGTTCTAGATGCGCATTGAT<br>AACAGGAGCTGTAAAAATGGCCCTATGCCC | for ADZ112 generation |
| prCE662 | AACAAATTTTACACATATTTGTGATCGCATTTTAATCCTG<br>GAGTCTGAAACATCATTTTTTGGATTATAT | for ADZ113 generation |
| prCE663 | TGCAGAAGTTGAACTCACTGTAACCAGGCCGTGTTGTTAG<br>GAAATGTATTTTATTAAATATTGCGGACAA | for ADZ114 generation |
| ***primers*** | | |
| prCE509 | CATGGTAAGTAGTTTCAGTTTTAAATGGAACAATTGAATA<br>TCCTTGCAGTCGGACACAATTCTTGGTTTCAACAAATC | for repair template for ADZ72 generation |
| prCE510 | GAATGAATGTGCAACAACTGAATGTGCAGAAAACTCCTT<br>TATCCGGAAGATCCTCCATCCAGC | |
| VL047 | GATATCTGGATCCACGAAGC | for cloning of pVL033 |
| VL048 | CTGCAGGAATTCCTCGAGAC | |
| VL049 | CGCACCGTACGTCTCGAGGAATTCCTGCAGGCATTCGAC<br>GAGATGAAGAA | |
| VL050 | ACCATGGGAAGCTTCGTGGATCCAGATATCCTATGTTCAG<br>TGCATTACGTTTTTG | |
| VL182 | ATAGCTTGGCGTAATCATGG | for cloning of pVL037, pVL038, pVL039, pVL040 and pVL073 |
| VL183 | ATAATTCACTGGCCGTCGTT | |
| VL183 | TATGACCATGATTACGCCAAGCTATCATCTCATGATGGGC<br>TACTTTC | |

| | | |
|---|---|---|
| VL184 | CCTTTGAGACCATGGTGTTGTATCAGATAAAACCTG | |
| VL185 | CTGATACAACACCATGGTCTCAAAGGGTGAAGAAG | |
| VL186 | AACATACACAAATCTACTTATACAATTCATCCATGCC | |
| VL187 | ATTGTATAAGTAGATTTGTGTATGTTGCTTCAGTC | |
| VL188 | TGTAAAACGACGGCCAGTGAATTATAACAGAAAAACTGT AGAATAGTTTTTTC | |
| VL189 | ATAGCTTGGCGTAATCATGG | |
| VL190 | CCTTTGAGACCATAAGATTATTGTAGAGTTTGTTGCAAC | |
| VL191 | CTACAATAATCTTATGGTCTCAAAGGGTGAAGAAG | |
| VL192 | TATGACCATGATTACGCCAAGCTATAGCCCACGCTGATCTT CTTC | |
| VL193 | CCTTTGAGACCATGTTTAGAATAGCCAGTTGTTGGAC | |
| VL194 | GGCTATTCTAAACATGGTCTCAAAGGGTGAAGAAG | |
| VL195 | ATCATATAGAGTACTACTTATACAATTCATCCATGCC | |
| VL196 | ATTGTATAAGTAGTACTCTATATGATCAACTCGAATAAAAA C | |
| VL197 | TGTAAAACGACGGCCAGTGAATTATGTTTAGACAAAAATT ATGTACTGTAAAAG | |
| VL198 | CCTTTGAGACCATATCGAGATTGAGGTAACGACG | |
| VL199 | CCTCAATCTCGATATGGTCTCAAAGGGTGAAGAAG | |
| VL378 | CTATGACCATGATTACGCCAAGCTATGATATTTTATGTGTCT CCATGATGATTATTTATAGT | |
| VL379 | TCACCCTTTGAGACCATAAGATTATTGTAGAGTTTGTTGCA ACCG | |
| VL380 | TGCAACAAACTCTACAATAATCTTATGGTCTCAAAGGGTG AAGAAGA | |
| VL381 | ACTGAAGCAACATATAGAACTCTACTTATACAATTCATCCA TGCCACC | |
| VL382 | GGATGAATTGTATAAGTAGAGTTCTATATGTTGCTTCAGTC GTTG | |
| VL383 | AACGACGGCCAGTGAATTATAAGTACATGGTTCTCCCATGT | |
| prCE576 | CTCGAGATTCAACTGATACCTCAACATG | for pCE090 cloning |
| prCE577 | TCTAGATGATGAAAACCTTCATTATATC | |
| prCE578 | CTCGAGTCTGATACAACACCATGTTC | for pCE091 cloning |
| prCE579 | TCTAGAGGAGTGCAACGACTGAAGCAAC | |
| prCE588 | GCTTCAGTCGTTGCACTCCTATTCAACTGATACCTCAAC | for pCE092 cloning |
| prCE589 | GTTTAGAATAGCCAGTTGTTGTGATGAAAACCTTCATTAT ATC | |
| prCE590 | GATATAATGAAGGTTTTCATCACAACAACTGGCTATTCTA AAC | |
| prCE591 | GGGAGACCGGCAGATCTGATCGAGTTGATCATATAGAG | |
| VL001 | TCAGGTTTCCACTGACAATG | *tent-5*; for genotyping |
| VL002 | TGATCTCGACCTGATATTCC | |
| VL003 | GTTCACACTCGTCCAACTC | |
| VL076 | ACGACAAGACAGGCGGTATC | *daf-16*; for genotyping |
| VL077 | TGAAGGGAGCCCATCAATGC | |
| VL078 | GGCAATCTGAGGTGATGATG | |
| prCE676 | GCATACTGTCGCTTCTTCATC | *daf-16::GFP*; for genotyping |
| prCE678 | TATTGCTGCTTACCTCACTCCTC | |
| prCE511 | GAATCTGATGTACGTGAATAGCGCAGTAG | in/active miniSOG; for genotyping |
| prCE512 | GCTCGACTACAGTGCATAAGATTTCTCAC | in/active miniSOG, *nspc-(8-10)*; for genotyping |
| prCE596 | GAGATCCTCGGCCGCAATGGT | active miniSOG; for genotyping |
| prCE597 | GAGATCCTCGGCCGCAATTGC | inactive miniSOG; for genotyping |

| | | |
|---|---|---|
| prCE649 | AAAACAACGAGTCGCTTAACAAAT | *nspc-(1-7)*; for genotyping |
| prCE650 | CTTCACCGGCCTTGAATTGC | |
| prCE609 | GAGTTTTGGGGCGTAGGATGA | |
| prCE651 | GACAACGTTACGACTCTGACC | *nspc-(8-10)*; for genotyping |
| prCE652 | GTAATGGACTTTGCGCCCTCT | |
| prCE654 | TGGCAAAAAGTCAGTTCAGGA | *nspc-(11-15)*; for genotyping |
| prCE655 | AAATTCTGAAAACAAAGTTCTGTGC | |
| prCE657 | GTAATATTTGTCTCGCTCAACTA | |
| prCE658 | ACGTGTTTCCACGTATTTTGTAG | *nspc-(16-20)*; for genotyping |
| prCE659 | TGAATTGAATGGTCATCCGGC | |
| prCE501 | TCACCAGTTGCTCACTCATTACAG | |
| prCE574 | ATGTTCCTTCGCACCCTCGTTG | *nspc-(1-15)*; for RT-qPCR |
| prCE575 | CTGATCCATGGAGAGCAGAATG | |
| prCE573 | ATGCTGGCTCTTCGCATCCTTG | *nspc-(16-20)*; for RT-qPCR |
| prCE547 | CTTGACAGAGGTTCAAGGTTATGG | |
